# Maternal sucrose consumption alters steroid levels in the mother, placenta, and fetus

**DOI:** 10.1101/2023.10.31.565065

**Authors:** Désirée R. Seib, Minseon M. Jung, Kiran K. Soma

**Author notes:** CORRESPONDING AUTHOR’S POSTAL AND EMAIL ADDRESS: Postal: Seib Lab, Duffy Science Centre, 550 University Ave, Charlottetown Prince Edward Island, C1A 4P3, Canada. Present address: Department of Biology University of Prince Edward Island, Charlottetown, PE, Canada.

## Abstract

Maternal diet has long-term effects on offspring brain development and behavior. Sucrose (table sugar) intakes are high in modern diets, but it is not clear how a maternal high-sucrose diet (HSD) affects the offspring. In rats, a maternal HSD (26% of calories from sucrose, which is human-relevant) alters maternal metabolism and brain and also alters adult offspring endocrinology and behavior in a sex-specific manner. Maternal sucrose intake increases corticosterone levels in adult female offspring and increases motivation for a sugar reward in adult male offspring. Here, to identify possible underlying mechanisms, we examined how a maternal HSD affects steroids in the dam, placenta, and fetus at embryonic day 19.5 using liquid chromatography tandem mass spectrometry. Maternal sucrose intake increased glucocorticoids (11-deoxycorticosterone and 11-dehydrocorticosterone) and tended to increase the mineralocorticoid aldosterone in maternal serum. In the placenta, maternal sucrose intake decreased androstenedione and testosterone. Maternal HSD increased aldosterone in the fetal blood. Similarly, in the fetal brain, maternal high-sucrose intake increased aldosterone in the medial prefrontal cortex and nucleus accumbens, decreased testosterone in the nucleus accumbens, and decreased corticosterone in the orbital cortex. In addition, the 11-dehydrocorticosterone/corticosterone and aldosterone/corticosterone ratios were increased in most examined brain regions. Lastly, maternal HSD increased 11-dehydrocorticosterone and aldosterone in the amniotic fluid. In summary, we found dramatic and widespread changes in maternal, placental, and fetal steroids that might mediate the long-term effects of maternal sucrose consumption on adult offspring neuroendocrinology and behavior.

## INTRODUCTION

The consumption of sucrose (table sugar) is high across the world (Newens & Walton 2016). Many individuals obtain 25% or more of their daily calories from sucrose and other added sugars (Ricciuto *et al*. 2021). There is evidence that high consumption of sucrose has negative health effects, such as increased risk for metabolic disorders and cardiovascular diseases (Stanhope 2016). Importantly, high sucrose intake also increases the risk for psychiatric disorders (Jacques *et al*. 2019) and has negative effects on cognitive function (Kendig 2014). In adult mice in a semi-natural environment, a high-sucrose diet (25% calories) reduces survival in females and reproduction in males (Ruff *et al*. 2013). In other experiments with adult rodents, high amounts of sucrose in the diet impair working memory, social memory, motivation and increase addictive behaviors (Wiss *et al*. 2017; Wong *et al*. 2017; Reichelt *et al*. 2019; Mizera *et al*. 2021). Thus, sucrose has negative effects on adult health and the brain.

There is very limited data on the effects of high-sucrose consumption during pregnancy on the mother and the developing fetal brain. Most rodent research on maternal diets focuses on high-fat diets (Naef *et al*. 2008; Vucetic *et al*. 2010) or high-fat/high-sugar diets (George *et al*. 2019). In these studies, there is an increase in maternal calorie intake and obesity, and it is difficult to dissociate the effects of the diet and the effects of maternal obesity (Vucetic *et al*. 2010). With high-fat/high-sugar diet studies, it is unclear if the observed effects are caused by an increase in fat or sugar. Moreover, in studies where sucrose is administered via drinking water, food intake is typically reduced, creating group differences (confounds) in protein, fat, vitamin, and mineral intakes (Hsu *et al*. 2015).

We have developed an animal model in which we feed rat dams a high-sucrose diet (HSD) that is isocaloric to the control (CON) diet, and micro- and macro-nutrients are matched (Tobiansky *et al*. 2020, 2021). The amount of sucrose (26% of calories) in the HSD is a human-relevant level, and the duration of exposure is relatively long (over 12 wk). Importantly, the HSD does not affect food consumption and body mass of rat dams (Tobiansky *et al*. 2020). Therefore, group differences can be unambiguously attributed to sucrose intake rather than to increased caloric intake or obesity or to decreased protein or fat intakes.

After approximately 16 wk, this HSD has negative effects on rat dams: impaired glucose tolerance, increased liver lipids, and increased transcript levels of *Emr1* (a macrophage marker) in gonadal adipose tissue. After weaning, HSD dams have decreased corticosterone levels in the blood but not in the brain (Tobiansky *et al*. 2020). All offspring were fed standard chow after weaning. In adult offspring, we observed long-lasting neural, endocrine, and behavioral effects of maternal HSD (Tobiansky *et al*. 2021). Maternal HSD increases baseline corticosterone levels in the blood and brain of female, but not male, offspring. Moreover, maternal HSD increases preference for a high-sucrose diet and a high-fat diet, increases motivation to work for sugar rewards in a progressive ratio task, and decreases mRNA levels of *Cyp17a1* (an androgenic enzyme) in the nucleus accumbens (NAc) of male, but not female, offspring (Tobiansky *et al*. 2021). These long-term effects of maternal HSD might be mediated by events during pregnancy.

The placenta is created during pregnancy and has two important functions. First, it is an essential source of steroids, such as allopregnanolone (Vacher *et al*. 2021). Second, the placenta protects the fetus from harmful substances in the mother’s bloodstream but allows nutrients and growth factors to pass to the fetus. For example, the placenta protects the fetus from high corticosterone levels in the mother by metabolizing corticosterone to 11-dehydrocorticosterone (DHC) via 11β-hydroxysteroid dehydrogenase 2 (11β-HSD2) (Cottrell *et al*. 2014).

In the present study, we use this model to identify potential mechanisms underlying the previously observed effects of maternal HSD on adult offspring. We focus on changes in steroid levels during pregnancy, particularly at embryonic day 19.5 (E19.5). The aim of our study was to determine the effects of a maternal HSD before and during gestation on the dam, placenta, and fetus. We focused on steroids in the dam’s serum, placenta (junctional zone and labyrinth zone), fetal blood, multiple microdissected fetal brain regions, and amniotic fluid. We focused on steroids in the dam’s serum, placenta (junctional zone and labyrinth zone), fetal blood, multiple microdissected fetal brain regions, and amniotic fluid. In the fetal brain, we examined the mesocorticolimbic system (orbital cortex (OC), medial prefrontal cortex (mPFC), nucleus accumbens (NAc), and ventral tegmental area (VTA)) as these brain regions are important for executive functions and motivated behaviours. In addition, we examined brain regions that are important for anxiety (amygdala (AMY)) and learning and memory (dorsal hippocampus (dHPC) and ventral hippocampus (vHPC)). Last, we examined the medial preoptic area (POA) and the hypothalamus (HYP) that modulate social behaviour and systemic hormones. All these brain regions are well-known to be impacted by early-life stress (Brunton 2015). We used liquid chromatography-tandem mass spectrometry (LC-MS/MS), which is highly specific and sensitive, to measure a panel of 11 steroids in very small tissue samples. We also monitored maternal weight, pregnancy outcomes, and fetal and placental weights.

## MATERIALS AND METHODS

### Animals and diets

All animal procedures were approved by the Animal Care Committee of the University of British Columbia and were in accordance with the Canadian Council on Animal Care. Animals were housed in the Centre for Disease Modeling at the University of British Columbia. Adult female Long-Evans rats (postnatal day 56-62, 125-175g, n=50, Charles River, Kingston, NY, USA) were pair housed in ventilated Allentown cages (22°C; 45 to 60% relative humidity) on a 12h:12h light/dark cycle (lights on at 10:30 am). Rats had *ad libitum* access to standard rat chow (Teklad Rodent Diet, catalog # 2918, Envigo, Madison, WI) for 1 wk after arrival. Rats had *ad libitum* access to water (purified by reverse osmosis and sterilized by chlorination) in plastic bottles throughout the experiment. We ran two cohorts of animals (cohort 1, n=30; cohort 2, n=20), and each week 5 CON and 5 HSD animals were mated.

We randomly assigned dams to either a control diet (CON) where 1% of calories were from sucrose, or a high-sucrose diet (HSD) where 26% of calories were from sucrose. The HSD contains a human-relevant level of sucrose intake (Ruff *et al*. 2013). One wk after arrival, the female rats were gradually introduced to either the CON diet (catalogue #: D12450ki; Research Diets, Inc., New Brunswick, NJ, USA) or a custom-made isocaloric, micro- and macro-nutrient matched HSD (catalogue #: D20061801I; Research Diets, Inc.; Table 1). In both diets, calories were derived from 20% protein, 70% carbohydrate, and 10% fat. In the HSD, 25.6% of calories were from sucrose and 30% of calories were from corn starch, whereas in the CON diet, 1% of calories were from sucrose (contained in Vitamin Mix) and 54% of calories were from corn starch. As a result, the starch content in the HSD was reduced, compared to the CON diet. CON and HSD females were on their respective diets for 10 wk prior to mating (Figure 1). Food consumption per cage and body mass were recorded weekly. Food consumption and body mass were not regularly measured after mating to avoid unnecessary stress to the pregnant females.

**Figure 1.**
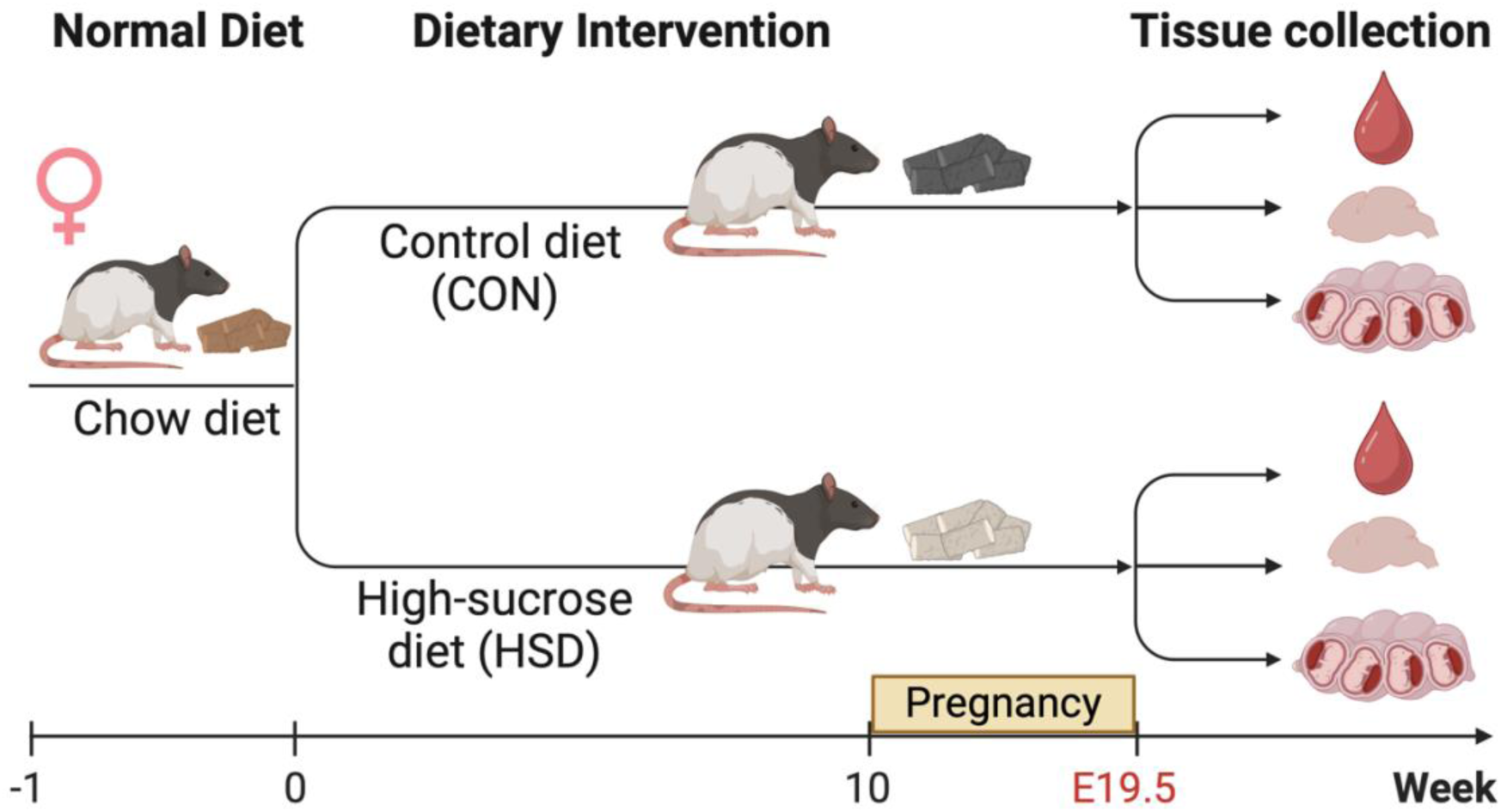
Experimental timeline. Adult female rats were fed a control (CON) diet or high-sucrose diet (HSD) for 10 wk. Then, females were mated. At embryonic day 19.5 (E19.5), maternal serum, fetal brains, placenta, amniotic fluid, and fetal blood were collected. Created with Biorender.com.

**Table 1.**
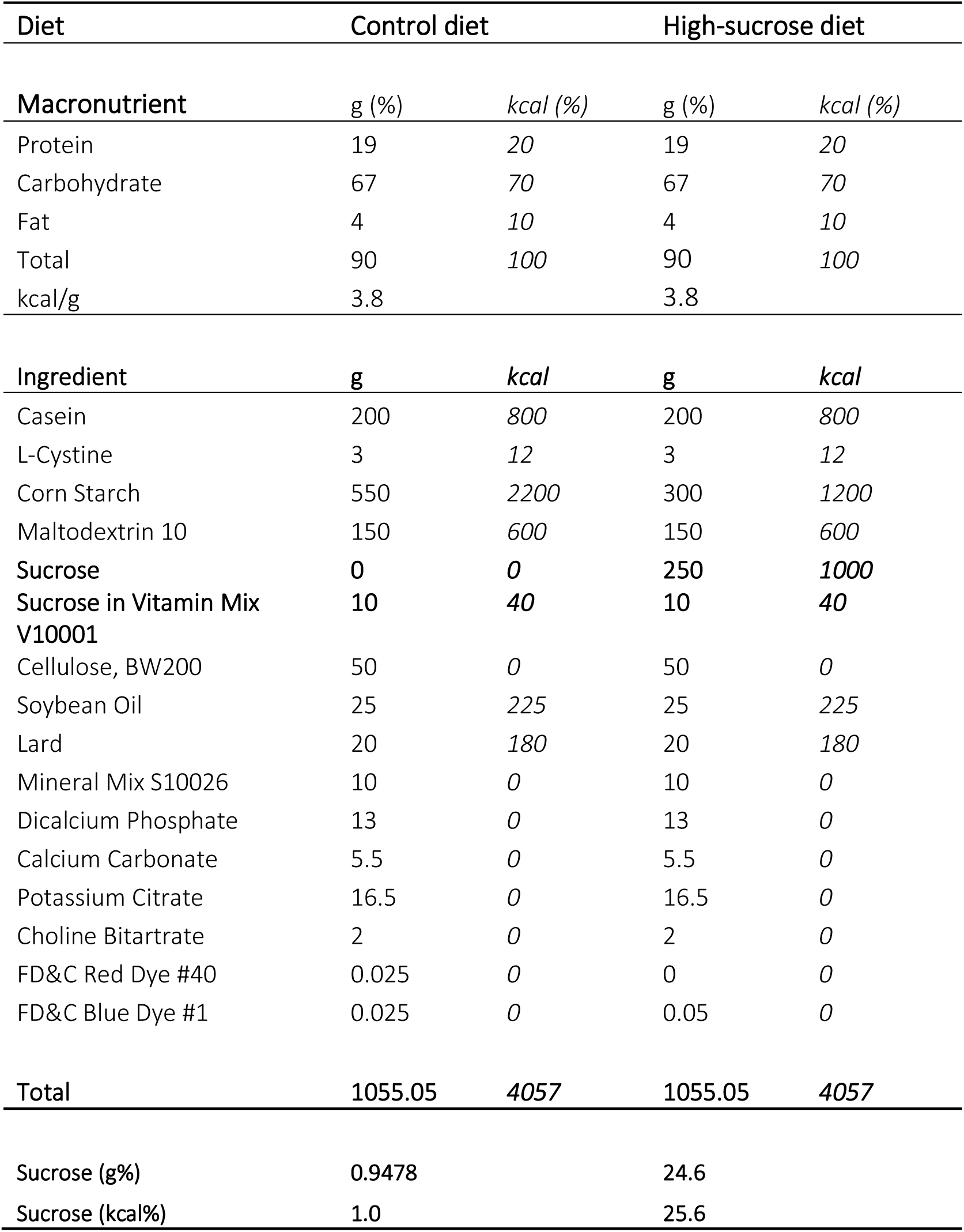
Diet composition.

| Diet | Control diet |  | High-sucrose diet |  |
| --- | --- | --- | --- | --- |
| Macronutrient | g (%) | kcal (%) | g (%) | kcal (%) |
| Protein | 19 | 20 | 19 | 20 |
| Carbohydrate | 67 | 70 | 67 | 70 |
| Fat | 4 | 10 | 4 | 10 |
| Total | 90 | 100 | 90 | 100 |
| kcal/g | 3.8 |  | 3.8 |  |
| Ingredient | g | kcal | g | kcal |
| Casein | 200 | 800 | 200 | 800 |
| L-Cystine | 3 | 12 | 3 | 12 |
| Corn Starch | 550 | 2200 | 300 | 1200 |
| Maltodextrin 10 | 150 | 600 | 150 | 600 |
| Sucrose | 0 | 0 | 250 | 1000 |
| Sucrose in Vitamin Mix V10001 | 10 | 40 | 10 | 40 |
| Cellulose, BW200 | 50 | 0 | 50 | 0 |
| Soybean Oil | 25 | 225 | 25 | 225 |
| Lard | 20 | 180 | 20 | 180 |
| Mineral Mix S10026 | 10 | 0 | 10 | 0 |
| Dicalcium Phosphate | 13 | 0 | 13 | 0 |
| Calcium Carbonate | 5.5 | 0 | 5.5 | 0 |
| Potassium Citrate | 16.5 | 0 | 16.5 | 0 |
| Choline Bitartrate | 2 | 0 | 2 | 0 |
| FD&C Red Dye #40 | 0.025 | 0 | 0 | 0 |
| FD&C Blue Dye #1 | 0.025 | 0 | 0.05 | 0 |
| Total | 1055.05 | 4057 | 1055.05 | 4057 |
| Sucrose (g%) | 0.9478 |  | 24.6 |  |
| Sucrose (kcal%) | 1.0 |  | 25.6 |  |

Adult male Long-Evans rats (postnatal day 56, n=50, Charles River, Kingston, NY, USA) were pair-housed for 2 wk upon arrival and always received standard rat chow except when housed with a female for mating. When the males were at least 10 wk old, each female was paired with a separate male. Vaginal lavages were collected from nulliparous females for estrous cycle staging. Once females entered proestrus or estrus, they were paired with a male in the evening. Vaginal lavage on the following morning was performed to detect sperm. If no sperm were detected and females had progressed in their cycle, then females were placed back in the home cages, and lavages were continued until females were in a receptive stage again. Then, mating was repeated as described before. If sperm were found, then this indicated embryonic day 0.5 (E0.5), the female was single-housed, and body mass was monitored to confirm the pregnancy until euthanasia at E19.5 (∼2 d before parturition).

### Tissue collection and tissue microdissection

On E19.5, 3-5h after lights on, rat dams were rapidly and deeply anesthetized using 5% isoflurane and rapidly decapitated. We chose E19.5 to avoid the increase in maternal glucocorticoids near parturition (E21.5) and to include the surge of testosterone in male fetuses from E16.5 to E20.5 (Martin *et al*. 1977; Ward *et al*. 2003). Trunk blood was collected within 2min of disturbance (1.77 ± 0.002 min) to avoid effects of stress on steroid levels (Taves *et al*. 2011). Whole blood was left on wet ice for 30 to 60 min and centrifuged at 5000g for 2min; then serum was collected and frozen on dry ice. The uterine horns were exposed, and the number of fetuses and the number of resorptions were determined. Fetuses were removed from the uterus within their amniotic sacs.

Amniotic fluid (20-100µl) was collected in a microcentrifuge tube by cutting the amniotic sac with fine scissors and then flash-frozen on dry ice. Then, the fetus and placenta were weighed to the nearest 1 mg. Subsequently, the fetus was decapitated and trunk blood (5-20µl) was collected using a P200 pipette. Up to eight fetuses per dam were collected and flash-frozen. For the steroid analysis, one male fetus and one female fetus per dam were used. If a dam only had male or female fetuses, then we used only one fetus per dam. Fetal blood, fetal brain (within the skull), and placenta were immediately flash frozen on dry ice and stored at -70°C. The tip of the tail of each fetus was collected and stored at -20°C for sex determination (Dhakal & Soares 2017).

### Steroid extraction and analysis

Fetal brains (within the skulls) and placenta were cryosectioned (300 µm thick) at -12 °C as before using M-1 embedding matrix (Thermo Scientific) (Tobiansky *et al*. 2018). Microdissection of brain regions and placenta was performed using the Palkovits punch technique with an Integra Miltex biopsy punch (1mm diameter; Fisher Scientific) (Palkovits 1973). Punch locations are shown in Figure S1 for fetal brain. Brain regions microdissected were the orbital cortex (OC), medial prefrontal cortex (mPFC), nucleus accumbens (NAc), amygdala (AMY), dorsal hippocampus (dHPC), ventral hippocampus (vHPC), medial preoptic area (POA), medial hypothalamus (HYP), and ventral tegmental area (VTA). Wet weight of tissue per punch for fetal brain is 0.199 mg, and, depending on the region, 2 to 12 punches in total were collected bilaterally. 20 punches were collected from the junctional zone, as well as from the labyrinth zone (Figure S2). Wet weight of tissue per punch for placenta is 0.183 mg. Weight of 1µl of maternal serum, fetal blood, and amniotic fluid is 1 mg.

We measured steroids in maternal serum (5µl), fetal blood (2µl), amniotic fluid (10µl), junctional zone (3.66mg), labyrinth zone (3.66mg), and fetal brain (0.4-2.4mg) as described in detail previously (Tobiansky *et al*. 2018; Hamden *et al*. 2021). Briefly, to each sample, 50µl of deuterated internal standards were added (Tobiansky *et al*. 2020; Hamden *et al*. 2021; Jalabert *et al*. 2021). Proteins were precipitated, and total steroids (bound and unbound) were extracted with 1ml HPLC-grade acetonitrile and homogenization in a bead mill homogenizer (Hamden *et al*. 2021). To remove neutral lipids, we added 1ml HPLC-grade hexane, vortexed and centrifuged samples, and then discarded the hexane. Samples were then dried in a vacuum centrifuge (60°C for 45 min). Steroids were resuspended in 55µl 25% HPLC-grade methanol:MilliQ water, centrifuged, and 50µl were transferred to a glass insert in an LC vial. Samples were stored at -20°C until analysis.

Blanks, quality controls, calibration curves, and samples were processed at the same time. Two multiple reaction monitoring transitions were used for each analyte, and one multiple reaction monitoring transition was used for each deuterated internal standard, as reported previously (Jalabert *et al*. 2021). Assay linearity, accuracy, precision, and matrix effects have been reported previously for pregnenolone, progesterone, corticosterone, 11-dehydrocorticosterone (DHC), androstenedione, testosterone, 17β-estradiol, and estrone (Tobiansky *et al*. 2018; Hamden *et al*. 2021; Jalabert *et al*. 2021). Maternal serum was analyzed in a second LC-MS/MS run for 11-deoxycorticosterone, using methods described before (Hamden *et al*. 2021). For allopregnanolone, aldosterone, and estriol, the transitions monitored and retention times were as follows (quantifier transition *m/z*; qualifier transition *m/z*; retention time): allopregnanolone (301.1 → 283.2; 301.1 → 135.0; 11.10min), aldosterone (361.3 → 343.3; 361.3 → 315.4; 2.83min), estriol (287.1 → 171.09; 287.1 → 144.9; 2.47min). Assay linearity, accuracy, precision, and matrix effects were validated for these steroids (Price *et al*. 2023). The lower limit of quantification (LLOQ) for progesterone, corticosterone, DHC, androstenedione, estrone, and estriol was 0.05pg. The LLOQ for testosterone and aldosterone was 0.03pg. The LLOQ for 17β-estradiol was 0.4pg. The LLOQ for pregnenolone and allopregnanolone was 2pg. All blanks were below the LLOQs, and all quality controls were acceptable.

### Data analyses

Chromatograms were analyzed using Multiquant. If an analyte was below the LLOQ, then that analyte was considered non-detectable. If detectable samples in a group were ≥ 40%, then non-detectable values were estimated via Gibbs sampler-based left-censored missing value imputation (GSIMP) using the METIMP 1.2 web tool (Wei *et al*. 2018a). Values missing at random were imputed using the random forest (RF) METIMP 1.2 web tool (Wei *et al*. 2018b). If less than 40% of samples in a group were detectable, then non-detectable values were set to LLOQ/SQRT(2) (Handelsman & Ly 2019).

When necessary, data were log-transformed prior to statistical analysis. Effects of maternal diet on steroids in maternal serum and gestational outcomes (not considering fetal sex) were analyzed by two-tailed independent t-tests. Effects of maternal diet and sex on fetal steroids and gestational outcomes (considering fetal sex) were analyzed using two-way ANOVA. A mixed model was used when values were missing. For all analyses, α was set at ≤ 0.05. All graphs are presented using non-transformed data (mean ± SEM). Statistical analyses were done in Prism 10 (GraphPad Software Inc., San Diego, CA, USA).

## RESULTS

### HSD did not affect dam food intake and body mass

Adult (12-wk-old) female rats were put on either a HSD (26% calories from sucrose) or an isocaloric and nutrient-matched CON diet (1% calories from sucrose; Table 1) for 10 wk before mating (Figure 1). The amount of food that the females consumed per cage was similar between groups (Figure 2A; diet: F_(1,23)_=0.55, p=0.47; time: F_(2.76,63.44)_=44.68, p<0.0001; diet x time: F_(9,207)_=0.60, p=0.79). Moreover, the body mass of CON and HSD females was similar (Figure 2B; diet: F_(1,48)_=0.01, p=0.91; time: F_(2.5,119.8)_=411.9, p<0.0001; diet x time: F_(10,480)_=0.72, p=0.71).

**Figure 2.**
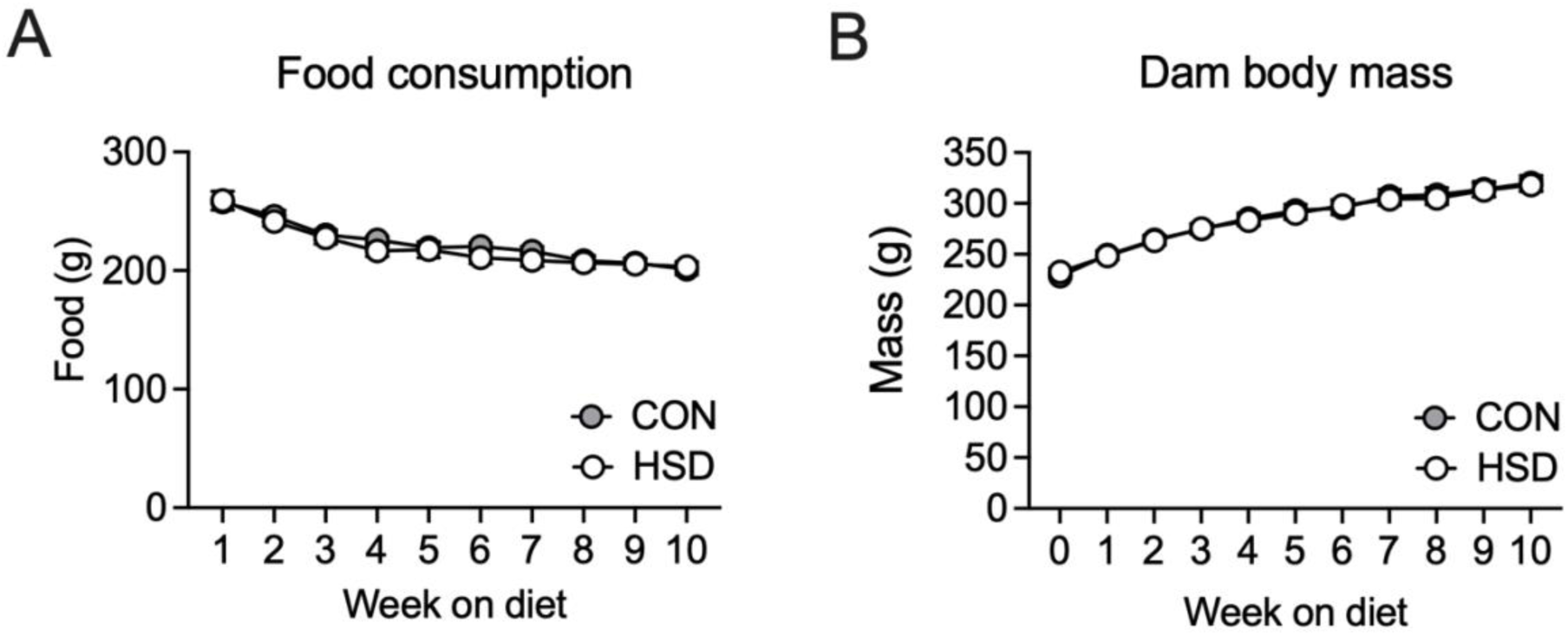
High-sucrose diet (HSD) does not alter food intake and body mass, relative to control (CON) diet. (**A**) Food intake per week and per cage of pair-housed CON diet and HSD females pre-gestation. (**B**) Female weight pre-gestation per week during the first 10 wk on the CON diet and HSD. n = 24-26.

### Maternal HSD reduced placental mass but did not affect other pregnancy outcomes

High-sucrose consumption before and during pregnancy did not significantly affect several pregnancy outcomes. Litter size tended to be higher in the HSD group (Figure 3A; *t*_(20)_=1.79, p=0.09). The numbers of male and female fetuses tended to be higher in the HSD group (Figure 3B; diet: F_(1,20)_=3.58, p=0.07; sex: F_(1,20)_=0.51, p=0.48; diet x sex: F_(1,20)_=0.51, p=0.48), and the percentage of male fetuses within a litter was not significantly different (Figure 3C; *t*_(20)_=1.5, p=0.15). The CON and HSD groups showed no difference in the number of resorptions (Figure 3D; *t*_(20)_=0.62, p=0.54). Maternal high-sucrose consumption did not affect the weight of the fetuses (Figure 3E; diet: F_(1,30)_=0.81, p=0.38; sex: F_(1,30)_=1.87, p=0.18; diet x sex: F_(1,30)_=0.14, p=0.71).

**Figure 3.**
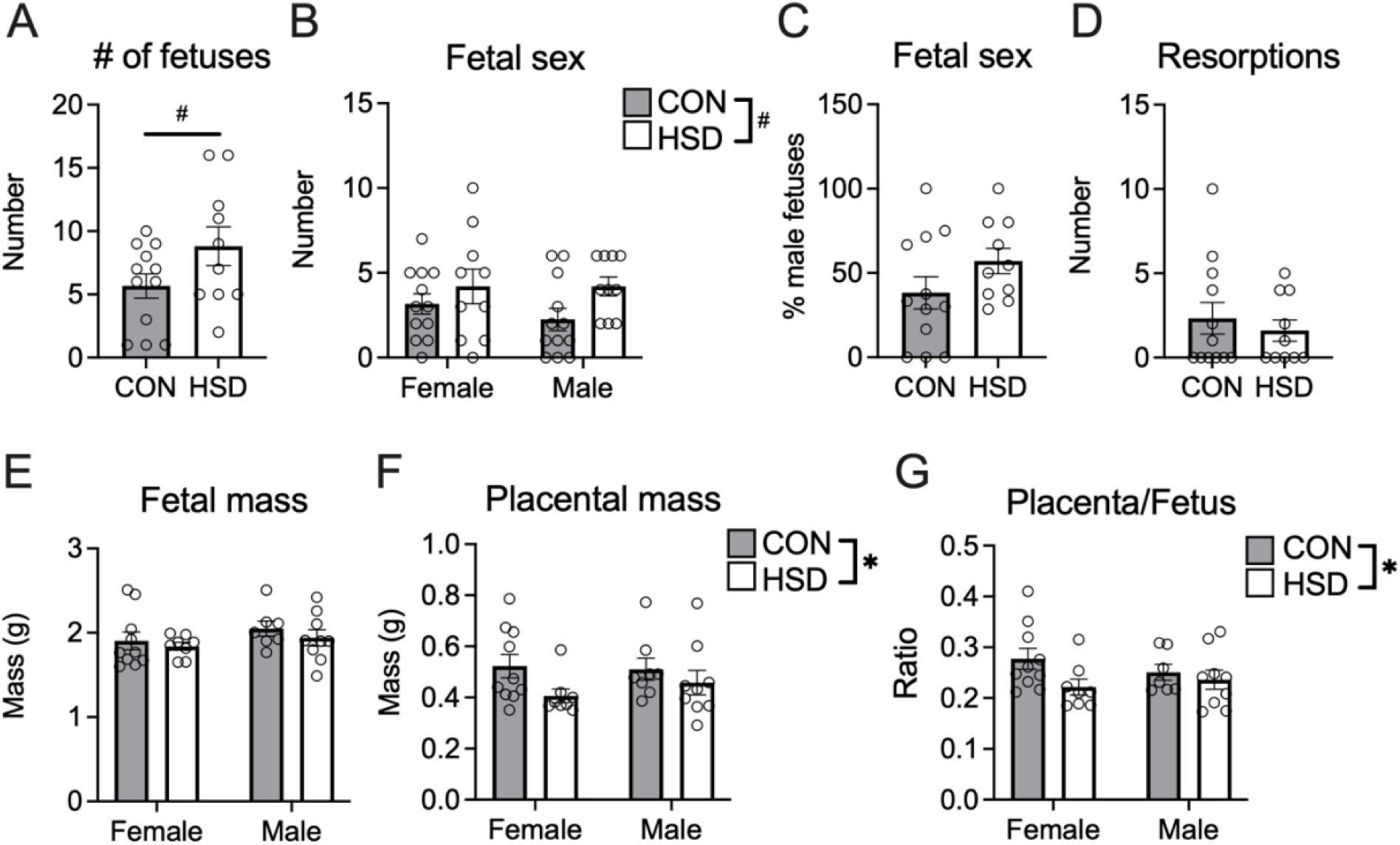
Maternal high sucrose consumption reduced placental mass. (**A**) The size of the litter tended to be increased in the HSD group as well as the (**B**) number of male and female fetuses (n=10-12). (**C**) The percentage of male fetuses was not affected by diet (n=10-12). (**D**) The number of resorptions (n = 10-12) and (**E**) fetal mass (n = 7-10) were similar between the CON and HSD groups. (**F**) Placental mass (n = 8-10) was reduced in the HSD group as well as the (**G**) placenta/fetus ratio (n = 7-10). CON, control; HSD, high-sucrose diet. ^#^p ≤ 0.1; *p ≤ 0.05.

However, the HSD significantly decreased placental mass (Figure 3F; diet: F_(1,31)_=4.75, p=0.04; sex: F_(1,31)_=0.26, p=0.62; diet x sex: F_(1,31)_=0.42, p=0.52) and the placenta/fetus mass ratio (Figure 3G; diet: F_(1,30)_=4.24, p=0.05; sex: F_(1,30)_=0.07, p=0.80; diet x sex: F_(1,30)_=1.03, p=0.32).

### Maternal HSD altered steroids in the maternal serum

We measured a panel of 11 steroids (pregnenolone, progesterone, allopregnanolone, corticosterone, DHC, aldosterone, androstenedione, testosterone, estrone, 17β-estradiol, estriol) in maternal serum at E19.5 (Figure 4). In the maternal serum, we found a trend for HSD to increase aldosterone levels (Figure 5A; *t*_(20)_=1.87, p=0.08). DHC levels in maternal serum were significantly increased in the HSD group (Figure 5B; *t*_(20)_=2.34, p=0.03). Levels of corticosterone (*t*_(20)_=0.91, p=0.38) and testosterone (*t*_(20)_=1.71, p=0.10) were not significantly different between CON and HSD dams (Figure 5C,D). Based on these data, we then measured 11-deoxycorticosterone (DOC) in maternal serum. DOC was significantly increased by HSD (Figure 5E; *t*_(18)_=2.1, p=0.05). Data for maternal steroids that were not significantly different between CON and HSD subjects are in Table S1.

**Figure 4.**
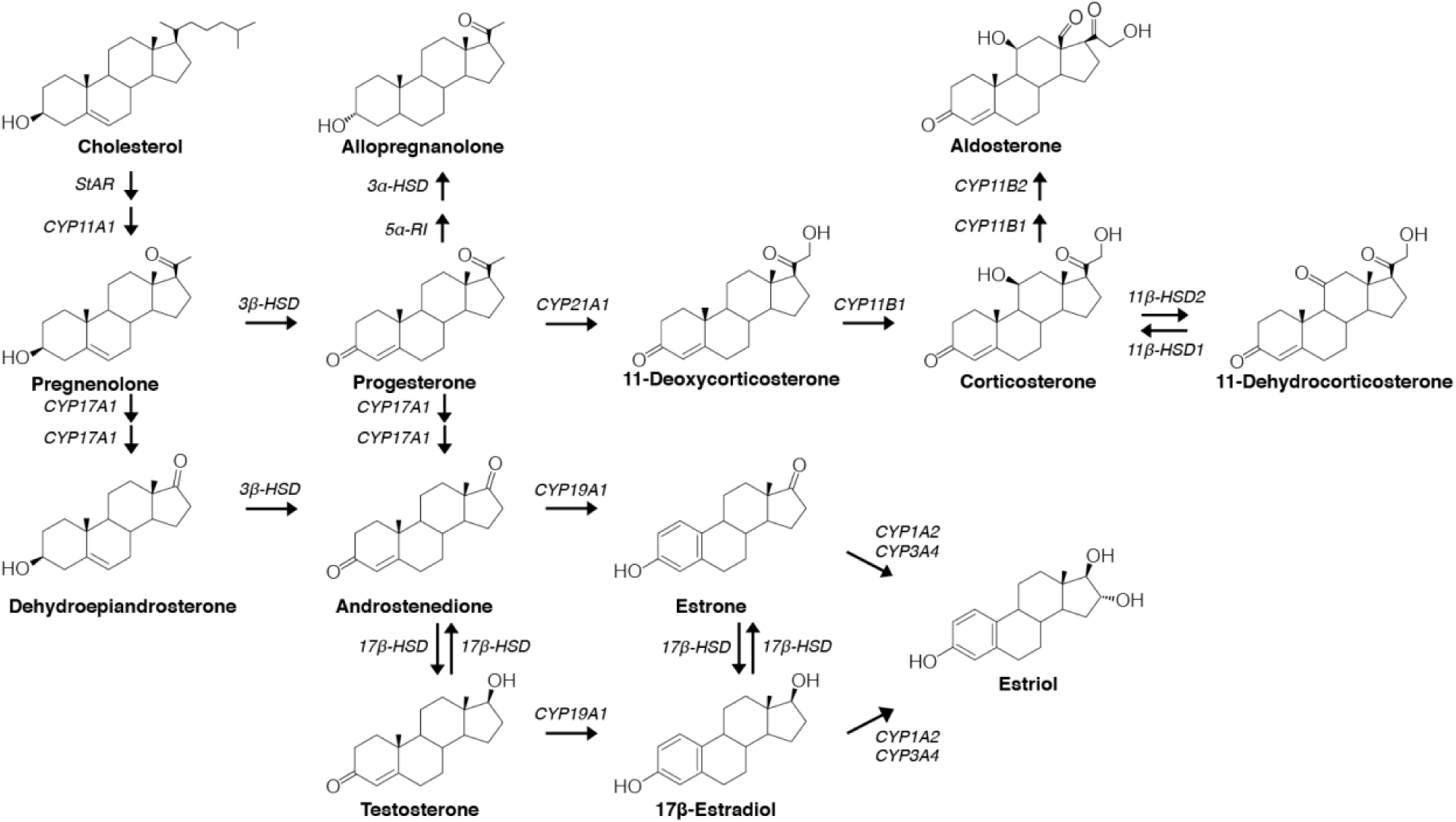
LC-MS/MS steroid panel. Overview of the steroidogenic pathway including the 11 steroids (pregnenolone, progesterone, allopregnanolone, corticosterone, DHC, aldosterone, androstenedione, testosterone, estrone, 17β-estradiol, estriol) that were analyzed with LC-MS/MS.

**Figure 5.**
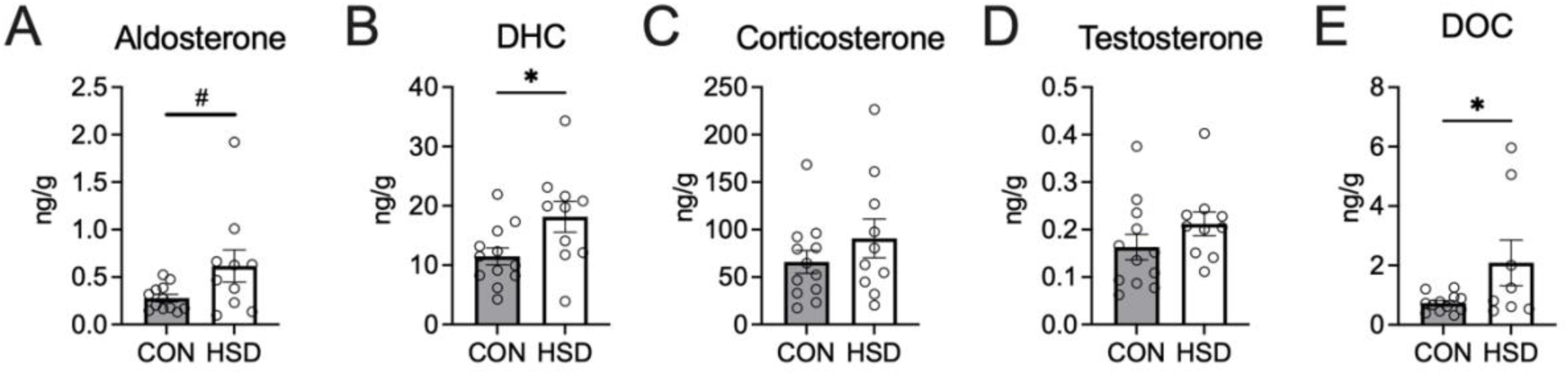
High sucrose intake alters maternal steroids. (**A**) HSD tended to increase aldosterone in maternal serum. (**B**) DHC was significantly increased by maternal HSD. (**C**) Corticosterone and (**D**) testosterone were not significantly affected by the HSD. (**E**) Maternal HSD significantly increased DOC in maternal serum. CON, control; HSD, high-sucrose diet; DHC, 11-dehydrocorticosterone; DOC, 11-deoxycorticosterone. n= 10-12. ^#^p ≤ 0.1; *p ≤ 0.05.

### Maternal HSD altered steroids in the placenta

The same 11 steroids were also measured in two parts of the placenta, the junctional zone and the labyrinth zone (Figure S2). HSD did not affect aldosterone in the junctional zone (Figure 6A; diet: F_(1,31)_=1.34, p=0.26; sex: F_(1,31)_=1.24, p=0.27; diet x sex: F_(1,31)_=0.03, p=0.86), but HSD significantly increased aldosterone in the labyrinth zone (Figure 6B; diet: F_(1,31)_=4.66, p=0.04; sex: F_(1,31)_=1.8, p=0.19; diet x sex: F_(1,31)_=0.62, p=0.44).

**Figure 6.**
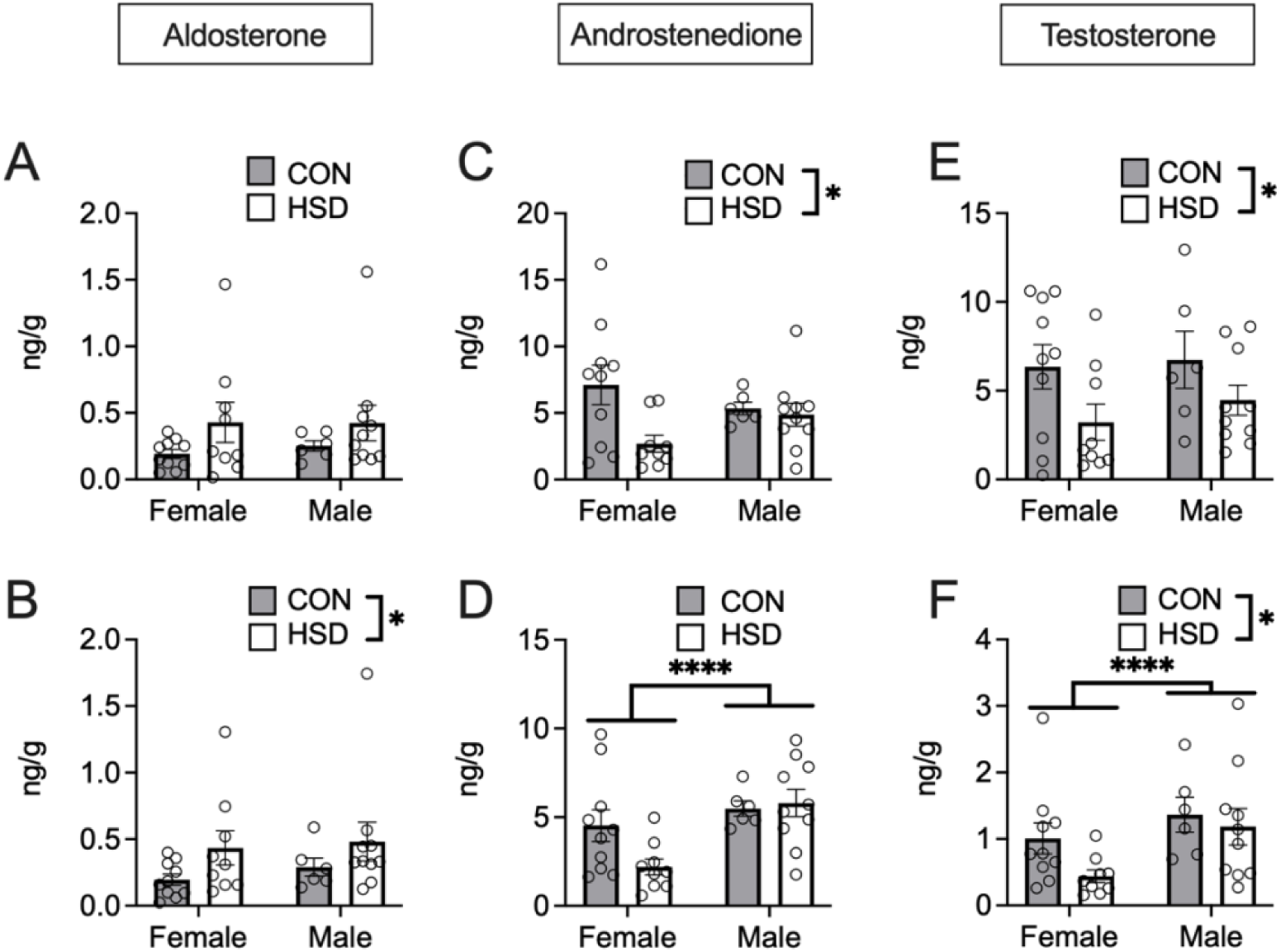
Placental steroids affected by maternal HSD. The upper row shows the results for the junctional zone and the lower row shows the results for the labyrinth zone. (**A**) Aldosterone in the junctional zone was not changed by the HSD. (**B**) Aldosterone in the labyrinth zone was increased by the HSD. (**C**) HSD significantly reduced androstenedione in the junctional zone. (**D**) Males had higher levels of androstenedione in the labyrinth zone than females, but levels were not altered by the HSD. (**E**) Diet significantly reduced testosterone in the junctional zone. (**F**) Males had significantly higher testosterone in the labyrinth zone than females, and the HSD significantly reduced testosterone. CON, control; HSD, high-sucrose diet. n=6-10. *, p ≤ 0.05; **, p ≤ 0.01

Androstenedione was significantly reduced by the HSD in the junctional zone (Figure 5C; diet: F_(1,31)_=5.12, p=0.03; sex: F_(1,31)_=0.04, p=0.85; diet x sex: F_(1,31)_=3.36, p=0.8) and was not affected by HSD in the labyrinth zone (Figure 6D; diet: F_(1,31)_=1.82, p=0.19; sex: F_(1,31)_=9.23, p<0.01; diet x sex: F_(1,31)_=3.12, p=0.09). Androstenedione levels were similar between males and females in the junctional zone (Figure 6C) but higher in males than in females in the labyrinth zone (Figure 6D). HSD did significantly reduce testosterone levels in the junctional zone, and testosterone levels were similar between males and females (Figure 6E; diet: F_(1,31)_=5.32, p=0.03; sex: F_(1,31)_=0.49, p=0.49; diet x sex: F_(1,31)_=0.14, p=0.72). Levels of testosterone were significantly higher in the labyrinth zone for males than females (Figure 6F; diet: F_(1,31)_=5.62, p=0.02; sex: F_(1,31)_=8.45, p<0.01; diet x sex: F_(1,31)_=1.17, p=0.29). HSD significantly reduced testosterone in the labyrinth zone.

DHC and corticosterone were not changed significantly in the (Table S2). Data from steroids in the placenta that remained unchanged by the maternal HSD are in Table S2.

### Maternal HSD altered steroids in the fetus

We also analyzed the same 11 steroids in fetal blood, fetal brain, and amniotic fluid.

### Fetal blood

Aldosterone was significantly increased by maternal HSD in fetal blood in both sexes (Figure 7A; diet: F_(1,35)_=20.27, p<0.0001; sex: F_(1,35)_=2.84, p=0.10; diet x sex: F_(1,35)_=0.69, p=0.41). DHC in fetal blood was not affected by the HSD or sex (Figure 7B; diet: F_(1,35)_=0.29, p=0.59; sex: F_(1,35)_=0.10, p=0.75; diet x sex: F_(1,35)_<0.01, p=0.98). Androstenedione (diet: F_(1,35)_=1.0, p=0.33; sex: F_(1,35)_=28.93, p<0.0001; diet x sex: F_(1,35)_=2.09, p=0.16) and testosterone (diet: F_(1,35)_=0.96, p=0.34; sex: F_(1,35)_=193.5, p>0.0001; diet x sex: F_(1,35)_=1.28, p=0.27) were not significantly altered by maternal HSD in the fetal blood (Figure 7C, D). As expected, both androgens in fetal blood were higher in males than in females. Results for steroids in fetal blood that were not affected by maternal diet are shown in Table S3.

**Figure 7.**
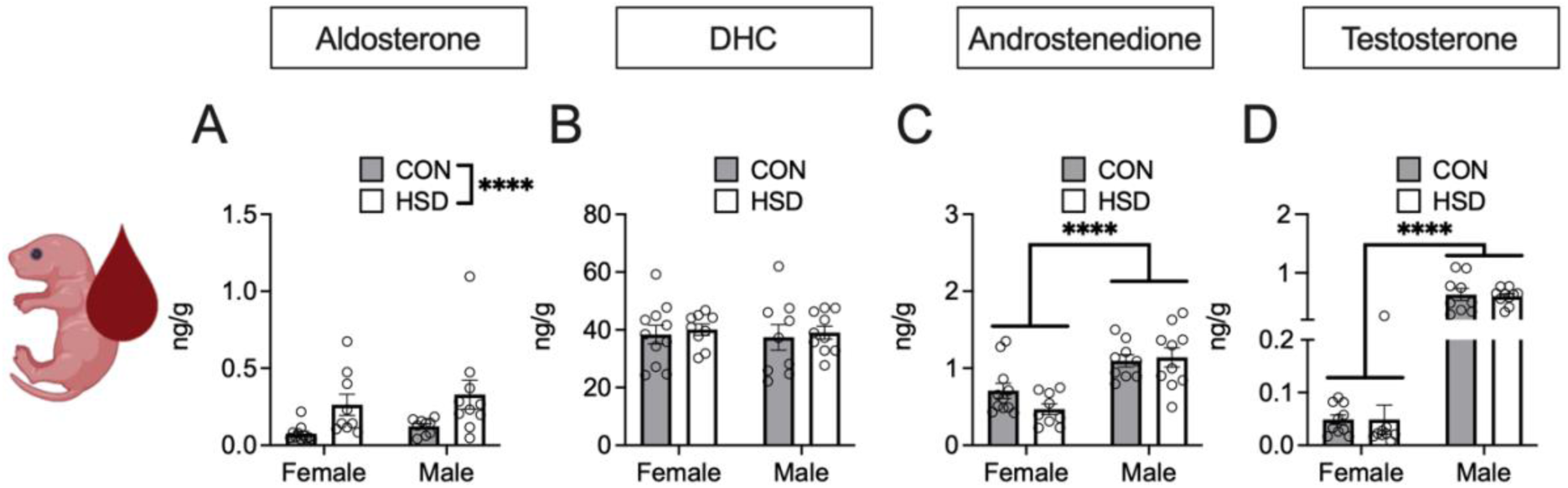
Chronic maternal high-sucrose intake alters steroids in fetal blood. (**A**) Aldosterone was significantly increased in the fetal blood due to maternal HSD. (**B**) DHC in fetal blood was not altered by maternal diet or sex. (**C**) Androstenedione in fetal blood was not altered due to maternal HSD but was significantly higher in males. (**D**) Testosterone in fetal blood was not affected by diet, but males had significantly higher testosterone than females. CON, control; HSD, high-sucrose diet; DHC, 11-dehydrocorticosterone. n=9-11; ***p ≤ 0.001; ****p ≤ 0.0001. Created with Biorender.com.

### Fetal brain

We analyzed 9 micro-dissected regions in the fetal brain and found region-specific effects of maternal HSD on androgens. Testosterone and androstenedione were not altered by maternal HSD in the OC (Figure 8A,B; testosterone: diet: F_(1,31)_=2.57, p=0.12; sex: F_(1,31)_=311.8, p<0.0001; diet x sex: F_(1,31)_=1.54, p=0.22; androstenedione: diet: F_(1,31)_=0.49, p=0.49; sex: F_(1,31)_=15.50, p<0.001; diet x sex: F_(1,31)_=1.81, p=0.19) and the mPFC (Figure 8C,D; testosterone: diet: F_(1,31)_=0.39, p=0.54; sex: F_(1,31)_=579.2, p<0.0001; diet x sex: F_(1,31)_=0.07, p=0.80; androstenedione: diet: F_(1,31)_=0.27, p=0.61; sex: F_(1,31)_=18.55, p<0.001; diet x sex: F_(1,31)_=0.48, p=0.50). But males had significantly higher levels of testosterone and androstenedione than females in both regions. In the NAc, testosterone was significantly reduced by maternal HSD (Figure 8E; diet: F(1,31)=4.50, p=0.04; sex: F_(1,31)_=241.9, p<0.0001; diet x sex: F_(1,31)_=3.82, p=0.06) and males had higher levels of testosterone than females. Androstenedione was not altered by maternal diet in the NAc (Figure 8F; diet: F_(1,31)_=0.52, p=0.48; sex: F_(1,31)_=19.30, p=0.0001; diet x sex: F_(1,31)_=1.51, p=0.23), and males had higher levels of androstenedione compared to females. Maternal HSD did not alter androgen levels in the POA, but males had higher levels of testosterone (Figure 8G; diet: F_(1,31)_=0.81, p=0.38; sex: F_(1,31)_=309.0, p<0.0001; diet x sex: F_(1,31)_<0.001, p=0.98) and androstenedione than females (Figure 8H; diet: F_(1,31)_=1.16, p=0.29; sex: F_(1,31)_=7.90, p<0.01; diet x sex: F_(1,31)_=0.12, p=0.73). Testosterone tended to be reduced by maternal HSD in the AMY (diet: F_(1,31)_=3.87, p=0.06; sex: F_(1,31)_=296.3, p<0.0001; diet x sex: F_(1,31)_=2.79, p=0.10; Table S4).

**Figure 8.**
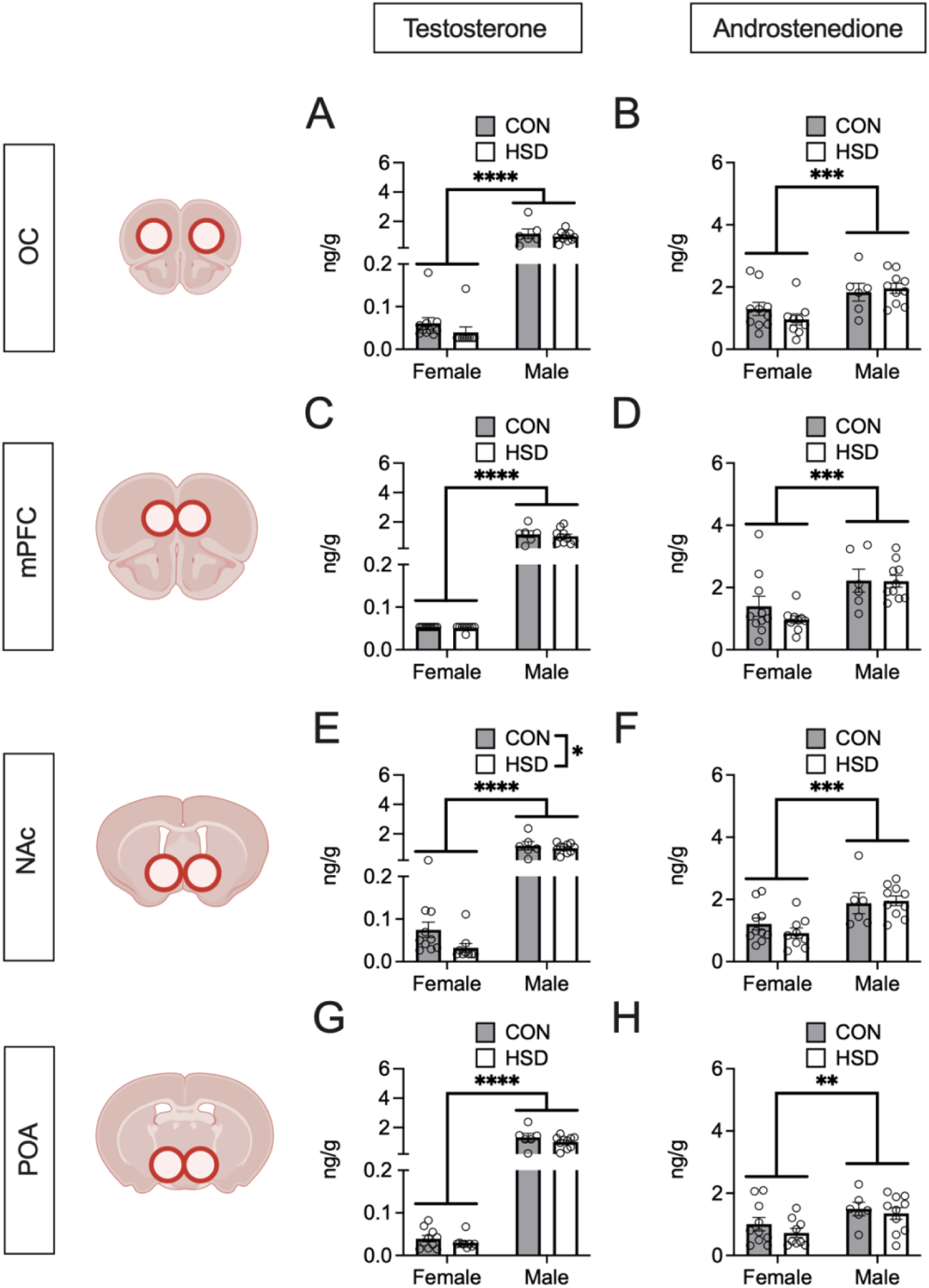
Maternal high-sucrose intake reduces testosterone in the nucleus accumbens. (**A**) Testosterone and (**B**) androstenedione were significantly higher in males compared to females in the OC and mPFC (C,D). Diet did not affect androgen levels in the OC and mPFC. Maternal high-sucrose intake significantly reduced testosterone (**E**) but not androstenedione (**F**) in the NAc. Males had significantly higher androgen levels than females in the NAc. (**G**) Testosterone and (**H**) androstenedione were significantly higher in males compared to females in the POA. Maternal diet did not alter androgen levels in the POA. OC, orbital cortex; mPFC, medial prefrontal cortex; NAc, nucleus accumbens; POA, preoptic area; CON, control; HSD, high-sucrose diet; n=6-10. *p ≤ 0.05; **p ≤ 0.01; ***p ≤ 0.001; ****p ≤ 0.0001. Created with Biorender.com.

We also found region-specific effects of maternal HSD on glucocorticoids and aldosterone. Corticosterone was significantly reduced by maternal HSD in the OC (Figure 9A; diet: F_(1,31)_=4.82, p=0.04; sex: F_(1,31)_<0.001, p=0.98; diet x sex: F_(1,31)_=0.24, p=0.63). DHC (Figure 9B; diet: F_(1,31)_=0.02, p=0.90; sex: F_(1,31)_<0.01, p=0.97; diet x sex: F_(1,31)_=3.06, p=0.09) and aldosterone (Figure 9C; diet: F_(1,31)_=2.25, p=0.14; sex: F_(1,31)_=1.36, p=0.25; diet x sex: F_(1,31)_=2.29, p=0.13) were not significantly altered by maternal HSD or sex in the OC. Corticosterone (Figure 9D; diet: F_(1,31)_=0.48, p=0.49; sex: F_(1,31)_=0.78, p=0.39; diet x sex: F_(1,31)_=0.71, p=0.40) and DHC (Figure 9E; diet: F_(1,31)_=0.12, p=0.74; sex: F_(1,31)_=0.18, p=0.67; diet x sex: F_(1,31)_=0.15, p=0.70) were not changed by maternal HSD or sex in the mPFC. Aldosterone was significantly increased by maternal HSD in the mPFC (Figure 9F; diet: F_(1,31)_=35.28, p<0.0001; sex: F_(1,31)_=2.83, p=0.10; diet x sex: F_(1,31)_=3.52, p=0.07). Maternal diet or sex had no effect on corticosterone (Figure 9G; diet: F_(1,31)_=1.60, p=0.22; sex: F_(1,31)_<0.001, p=0.98; diet x sex: F_(1,31)_=0.47, p=0.50) and DHC (Figure 9H; diet: F_(1,31)_=0.30, p=0.59; sex: F_(1,31)_=0.29, p=0.59; diet x sex: F(1,31)=1.32, p=0.26) in the NAc. Aldosterone was significantly increased by maternal HSD in the NAc (Figure 9I, diet: F_(1,31)_=5.46, p=0.03; sex: F_(1,31)_=0.09, p=0.76; diet x sex: F_(1,31)_<0.01, p=0.97). Corticosterone tended to be reduced in the POA by maternal HSD (Figure 9J, diet: F_(1,31)_=3.65, p=0.07; sex: F_(1,31)_=0.02, p=0.89; diet x sex: F_(1,31)_=1.94, p=0.17). DHC (Figure 9K, diet: F_(1,31)_=0.02, p=0.88; sex: F_(1,31)_=0.25, p=0.62; diet x sex: F_(1,31)_=0.01, p=0.92) and aldosterone (Figure 9L, diet: F_(1,31)_=2.64, p=0.11; sex: F_(1,31)_=0.04, p=0.84; diet x sex: F_(1,31)_=0.18, p=0.68) were not changed by maternal diet in the POA. Aldosterone showed a trend to be increased by maternal HSD in the AMY (diet: F_(1,31)_=3.11, p=0.09; sex: F_(1,31)_=1.53, p=0.23; diet x sex: F_(1,31)_=1.54, p=0.22; Table S4). All steroid data for the fetal brain regions are presented in Table S4.

**Figure 9.**
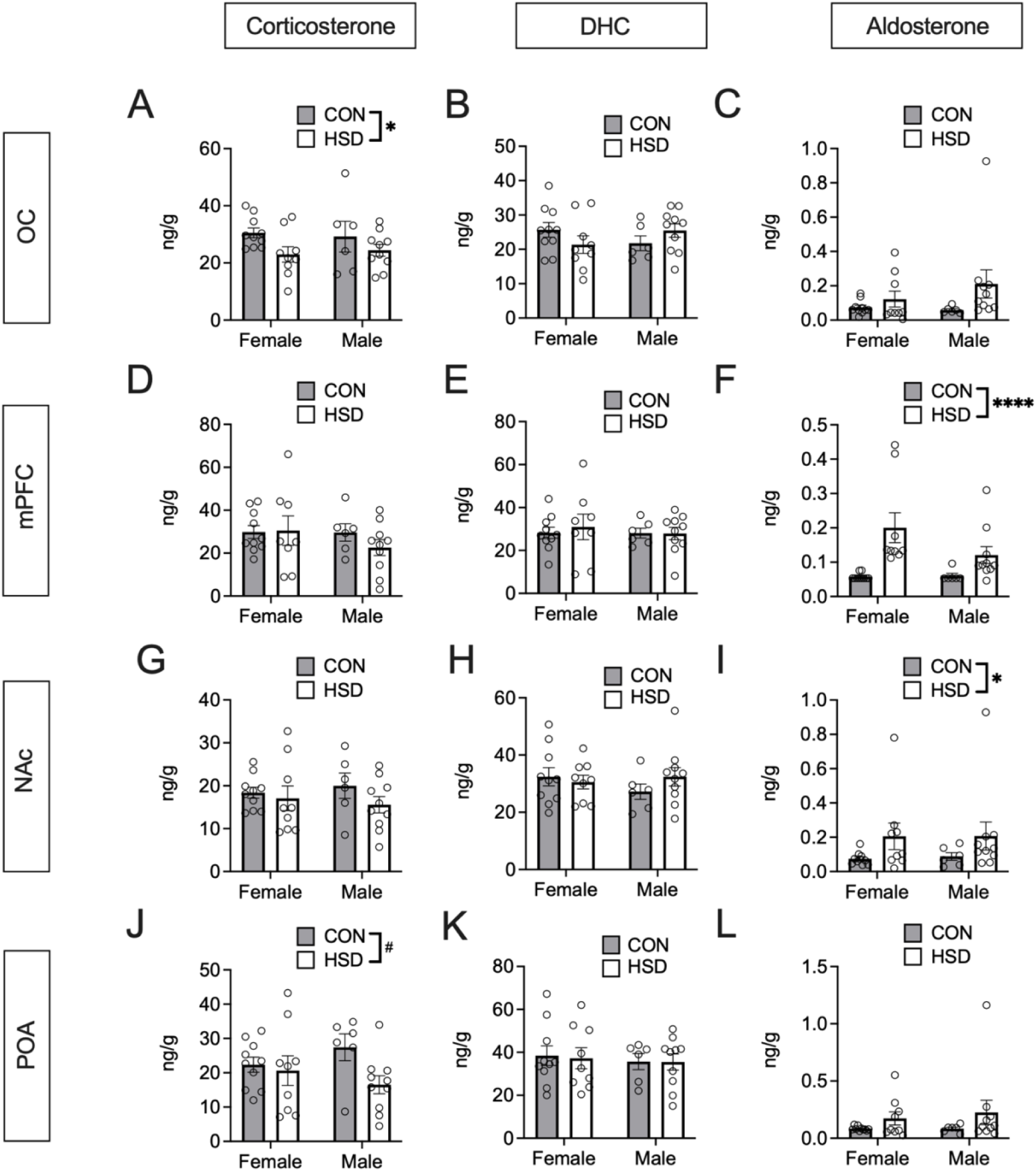
Chronic maternal high-sucrose intake alters fetal steroids in the brain. (**A**) Corticosterone was significantly reduced in the OC by maternal HSD. (**B**) DHC and aldosterone (**C**) were not significantly altered by maternal diet in the OC. (**D**) Maternal high-sucrose intake did not alter corticosterone or (**E**) DHC in the fetal mPFC. (**F**) Aldosterone in fetal NAc was significantly increased due to maternal HSD. (**G**) Corticosterone and DHC (**H**) levels in the NAc were not altered by maternal diet. (**I**) Aldosterone was significantly increased by maternal HSD in the fetal NAc. (**J**) Corticosterone tended to be reduced in the fetal preoptic area by maternal HSD. (**K**) DHC and aldosterone (**L**) were not significantly altered by maternal HSD in the preoptic area. OC, orbital cortex; mPFC, medial prefrontal cortex; NAc, nucleus accumbens; POA, preoptic area; CON, control; HSD, high-sucrose diet; DHC, 11-dehydrocorticosterone; n=6-10. ^#^p ≤ 0.1; *p ≤ 0.05; ****p ≤ 0.0001.

As in previous studies (Vagnerová *et al*. 2023), we also analyzed the ratios of DHC/corticosterone and aldosterone/corticosterone, as corticosterone is the precursor to DHC and aldosterone. Interestingly, both ratios were increased by maternal HSD in most brain regions. In the OC, DHC/corticosterone tended to be increased (Table 2; diet: F_(1,34)_=3.76, p=0.06; sex: F_(1,34)_=0.13, p=0.72; diet x sex: F_(1,34)_=0.22, p=0.64) and aldosterone/corticosterone (Table 2; diet: F_(1,34)_=4.83, p=0.04; sex: F_(1,34)_=1.38, p=0.25; diet x sex: F_(1,34)_=1.39, p=0.25) was significantly increased by maternal HSD. Importantly, the mPFC was the only examined brain region where maternal HSD did not increase or tend to increase the DHC/corticosterone ratio (Table 2; diet: F_(1,34)_=1.83, p=0.19; sex: F_(1,34)_=0.32, p=0.58; diet x sex: F_(1,34)_=0.06, p=0.81). However, the aldosterone/corticosterone ratio was significantly increased by maternal HSD in the mPFC (Table 2; diet: F_(1,34)_=26.53, p<0.0001; sex: F_(1,34)_=0.20, p=0.66; diet x sex: F_(1,34)_=0.37, p=0.55). In the NAc, both DHC/corticosterone (Table 2; diet: F_(1,34)_=6.55, p=0.02; sex: F_(1,34)_<0.01, p=0.96; diet x sex: F_(1,34)_=1.31, p=0.26) and aldosterone/corticosterone (Table 2; diet: F_(1,34)_=10.95, p<0.01; sex: F(1,34)=0.15, p=0.71; diet x sex: F(1,34)=0.06, p=0.81) were significantly increased by maternal HSD. Similarly, both DHC/corticosterone (Table 2; diet: F_(1,34)_=7.67, p<0.01; sex: F_(1,34)_<0.01, p=0.94; diet x sex: F_(1,34)_=1.54, p=0.22) and aldosterone/corticosterone (Table 2; diet: F_(1,34)_=9.09, p<0.01; sex: F_(1,34)_=0.04, p=0.84; diet x sex: F_(1,34)_=1.06, p=0.31) were significantly increased in the POA by maternal HSD. Ratios for all brain regions are presented in Table 2.

**Table 2.** DHC/corticosterone and aldosterone/corticosterone ratios in fetal brain regions.

|  | CON Female |  |  | HSD Female |  |  | CON Male |  |  | HSD Male |  |  |
| --- | --- | --- | --- | --- | --- | --- | --- | --- | --- | --- | --- | --- |
| *DHC/CORT* |  |  |  |  |  |  |  |  |  |  |  |  |
| OC | 0.85 | ± | 0.06 | 0.99 | ± | 0.13 | 0.84 | ± | 0.12 | 1.07 | ± | 0.08 |
| mPFC | 0.98 | ± | 0.09 | 1.07 | ± | 0.06 | 1.02 | ± | 0.11 | 2.05 | ± | 0.88 |
| NAc | **1.76** | ± | **0.11** | **2.05** | ± | **0.23** | **1.51** | ± | **0.23** | **2.28** | ± | **0.23** |
| AMY | **1.26** | ± | **0.07** | **1.58** | ± | **0.16** | **1.23** | ± | **0.15** | **1.60** | ± | **0.14** |
| POA | **0.61** | ± | **0.05** | **0.52** | ± | **0.06** | **0.77** | ± | **0.13** | **0.46** | ± | **0.06** |
| dHPC | **0.81** | ± | **0.05** | **1.04** | ± | **0.12** | **0.77** | ± | **0.06** | **0.98** | ± | **0.09** |
| vHPC | **0.99** | ± | **0.07** | **1.32** | ± | **0.09** | **1.04** | ± | **0.16** | **1.18** | ± | **0.10** |
| HYP | **0.95** | ± | **0.07** | **1.20** | ± | **0.06** | **1.00** | ± | **0.12** | **1.35** | ± | **0.10** |
| VTA | **1.10** | ± | **0.09** | **1.25** | ± | **0.11** | **0.86** | ± | **0.11** | **1.24** | ± | **0.11** |
| *Aldosterone/CORT* |  |  |  |  |  |  |  |  |  |  |  |  |
| OC | 0.002 | ± | 0.000 | 0.005 | ± | 0.002 | 0.003 | ± | 0.001 | 0.010 | ± | 0.004 |
| mPFC | 0.002 | ± | 0.000 | 0.008 | ± | 0.001 | 0.002 | ± | 0.000 | 0.009 | ± | 0.003 |
| NAc | 0.004 | ± | 0.000 | 0.010 | ± | 0.002 | 0.006 | ± | 0.003 | 0.015 | ± | 0.005 |
| AMY | 0.003 | ± | 0.000 | 0.009 | ± | 0.002 | 0.007 | ± | 0.002 | 0.011 | ± | 0.004 |
| POA | 0.004 | ± | 0.000 | 0.008 | ± | 0.002 | 0.004 | ± | 0.001 | 0.016 | ± | 0.007 |
| dHPC | 0.003 | ± | 0.001 | 0.006 | ± | 0.002 | 0.004 | ± | 0.001 | 0.009 | ± | 0.003 |
| vHPC | 0.003 | ± | 0.001 | 0.007 | ± | 0.002 | 0.005 | ± | 0.003 | 0.011 | ± | 0.004 |
| HYP | 0.003 | ± | 0.001 | 0.007 | ± | 0.002 | 0.005 | ± | 0.002 | 0.012 | ± | 0.005 |
| VTA | 0.003 | ± | 0.000 | 0.008 | ± | 0.002 | 0.004 | ± | 0.002 | 0.013 | ± | 0.005 |

Data are shown as mean ± s.e.m. n=6-10. Two-way ANOVA was used for statistical analysis to examine the effects of diet and sex on different brain regions. Main effect of diet: data shown in **bold**, $p \leq 0.05$ . OC, orbital cortex; mPFC, medial prefrontal cortex; NAc, nucleus accumbens; AMY, amygdala; POA, preoptic area; dHPC, dorsal hippocampus; vHPC, ventral hippocampus; HYP, hypothalamus; VTA, ventral tegmental area; CON, control; HSD, high-sucrose diet; DHC, 11-dehydrocorticosterone; CORT, corticosterone.

### Amniotic fluid

Aldosterone was significantly increased by HSD in the amniotic fluid of males and females (Figure 10A; diet: F_(1,34)_=5.60, p=0.02; sex: F_(1,34)_=0.59, p=0.45; diet x sex: F_(1,34)_=0.24, p=0.63). DHC was also significantly increased by HSD in the amniotic fluid in both sexes (Figure 10B; diet: F_(1,34)_=5.04, p=0.03; sex: F_(1,34)_=0.40, p=0.53; diet x sex: F_(1,34)_<0.001, p=0.99). Androgens in the amniotic fluid were not altered by maternal diet, but androstenedione (Figure 10C; diet: F_(1,34)_=0.85, p=0.36; sex: F_(1,34)_=23.98, p<0.0001; diet x sex: F_(1,34)_=0.34, p=0.56) and testosterone (Figure 10D; diet: F_(1,34)_=0.05, p=0.82; sex: F_(1,34)_=184.5, p<0.0001; diet x sex F_(1,34)_=0.59, p=0.45) were significantly higher in males than females. Other steroid levels in the amniotic fluid are presented in Table S3.

**Figure 10.**
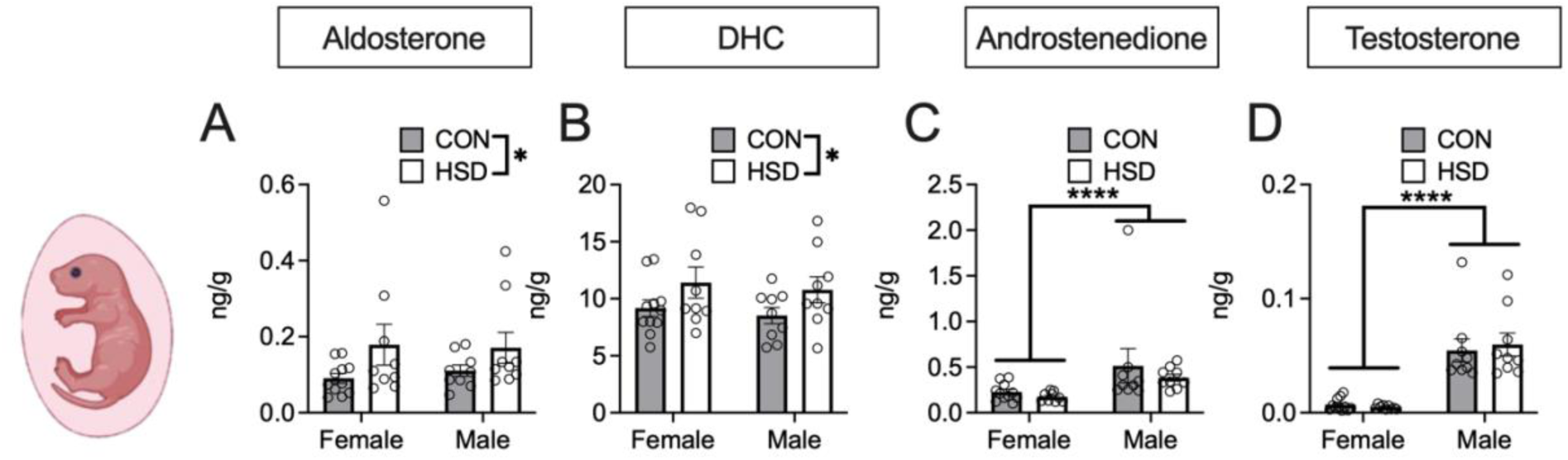
Chronic maternal high-sucrose intake alters steroids in the amniotic fluid. Aldosterone (**A**) and DHC (**B**) were significantly increased in the amniotic fluid due to maternal HSD. Androstenedione (**C**) and testosterone (**D**) were significantly higher in male than female fetuses but not affected by maternal diet. CON, control; HSD, high-sucrose diet; DHC, 11-dehydrocorticosterone. n=9-11. *p ≤ 0.05; ****p ≤ 0.0001. Created with Biorender.com.

## DISCUSSION

In this study, we investigated potential underlying mechanisms of the long-term endocrine and behavioral effects that a maternal HSD had on the adult offspring (Tobiansky *et al*. 2021). Our HSD contains a human-relevant level of sucrose (26% of kcal) and was provided over an extended period (10 wk prior to and during gestation) to mimic long-term human exposure. The control diet contained 1% kcal from sucrose and was isocaloric and micronutrient- and macronutrient-matched. Starch content was reduced in the HSD compared to the CON diet. Dam food intake and body mass were not affected by the HSD, as before (Tobiansky *et al*. 2020). Thus, this is not an obesity model, and the effects of sucrose intake cannot be attributed to increased maternal weight. This novel animal model was combined with microdissection of the placenta and fetal brain and highly sensitive and specific LC-MS/MS assays of 11 steroids.

Maternal HSD had effects on multiple steroids and multiple tissues. Steroid levels in a specific tissue can be determined by several factors, such as steroid levels in the blood, steroid binding globulins, local synthesis and metabolism, diffusion into the blood, local sequestration, and local receptor binding. Maternal HSD significantly increased DOC and DHC in maternal serum. Maternal sucrose intake also significantly reduced placenta mass, significantly increased aldosterone in the labyrinth zone, significantly decreased androstenedione in the junctional zone, and significantly reduced testosterone in both regions of the placenta (Table 3). In fetal blood, aldosterone was significantly increased by maternal HSD. In the fetal brain, maternal HSD significantly reduced testosterone in the NAc, significantly increased aldosterone in the mPFC and NAc, significantly decreased corticosterone in the OC, and significantly increased DHC/corticosterone and aldosterone/corticosterone ratios in almost all examined regions (Table 4). In the amniotic fluid, maternal HSD significantly increased DHC and aldosterone (Table 3).

**Table 3.** Summary of steroid data in maternal and fetal tissues.

|  | Maternal serum | Junctional zone | Labyrinth zone | Fetal blood | Amniotic fluid |
| --- | --- | --- | --- | --- | --- |
| Testosterone | - | ↓ | ↓ | - | - |
| Androstenedione | - | ↓ | - | - | - |
| 11-Dehydrocorticosterone | ↑ | - | - | - | ↑ |
| Aldosterone | - | - | ↑ | ↑ | ↑ |
| Corticosterone | - | - | - | - | - |
| DHC/corticosterone | - | - | - | - | - |
| Aldosterone/corticosterone | - | ↑ | ↑ | ↑ | ↑ |

Arrows indicate a significant increase or decrease in steroid concentration caused by maternal high-sucrose diet. Dashes (–) indicate no significant effect of maternal high-sucrose diet.

**Table 4.** Summary of steroid data in fetal brain.

|  | OC | mPFC | NAc | AMY | POA | dHPC | vHPC | HYP | VTA |
| --- | --- | --- | --- | --- | --- | --- | --- | --- | --- |
| Testosterone | - | - | ↓ | - | - | - | - | - | - |
| Androstenedione | - | - | - | - | - | - | - | - | - |
| 11-Dehydrocorticosterone | - | - | - | - | - | - | - | - | - |
| Aldosterone | - | ↑ | ↑ | - | - | - | - | - | - |
| Corticosterone | ↓ | - | - | - | - | - | - | - | - |
| DHC/corticosterone | - | - | ↑ | ↑ | ↓ | ↑ | ↑ | ↑ | ↑ |
| Aldosterone/corticosterone | ↑ | ↑ | ↑ | ↑ | ↑ | - | ↑ | - | ↑ |

Arrows indicate a significant increase or decrease caused by maternal high-sucrose diet. Dashes (–) indicate no significant effect of maternal high-sucrose diet.
OC, orbital cortex; mPFC, medial prefrontal cortex; NAc, nucleus accumbens; AMY, amygdala; POA, preoptic area; dHPC, dorsal hippocampus; vHPC, ventral hippocampus; HYP, hypothalamus; VTA, ventral tegmental area.

Taken together, these data suggest that high sucrose intake is a stressor that increases glucocorticoid and aldosterone signaling and reduces androgen signaling, which might account for the enduring effects of maternal HSD on offspring hormones, brain, and behavior.

### Maternal HSD alters glucocorticoid levels

In the dam’s serum, the HSD increased glucocorticoids. The corticosterone precursor DOC and the corticosterone metabolite DHC were significantly increased in maternal serum, but interestingly, corticosterone itself was not altered significantly. While DOC and DHC levels are elevated in the maternal circulation, the dams maintain normal corticosterone levels, perhaps because high corticosterone levels are harmful to the fetus (Cottrell *et al*. 2014; Brunton & Walker 2024). This might be achieved, in part, via increased hepatic metabolism of corticosterone to DHC and other metabolites. Another study used LC-MS/MS to measure glucocorticoid levels in response to dietary changes in pregnant rat dams and fetuses. In the dams, the authors observed similar levels of DHC and progesterone but 2-3 times higher corticosterone levels (Crew *et al*. 2016). Similarly, in the fetuses, Crew et al. observed slightly higher levels of circulating corticosterone. These data suggest a strain difference (Wistar rats vs. Long Evans rats) or a timing difference (E15 and E21 vs. E19.5).

Glucocorticoids were affected differently in the fetal brain and the amniotic fluid. In the fetal brain, corticosterone was reduced in the OC and tended to be reduced in the POA by maternal HSD. Both of those regions act on the hypothalamus to program the HPA axis during development (Viau & Meaney 1996; Sullivan & Gratton 2002; Smith & Vale 2006; Herman *et al*. 2020). The OC gives rise to prefrontal regions that play important roles in executive functions and decision making (Klein-Flügge *et al*. 2022). The POA expresses glucocorticoid receptors and thus is able to respond to glucocorticoid changes (Briski *et al*. 1997). The POA is well-known for its role in social behaviors, including male reproductive behavior and female parental behavior (Tsuneoka & Funato 2021). Remarkably, when we examined DHC/corticosterone and aldosterone/corticosterone ratios, these ratios were elevated significantly in almost all brain regions of male and female fetuses. This suggests that the fetus can maintain corticosterone at normal levels via metabolism to DHC, aldosterone, or other metabolites.

DHC but not corticosterone was also increased in the amniotic fluid by maternal HSD. Over time though, the offspring might not be able to maintain normal corticosterone levels, as we saw increased baseline blood corticosterone levels in adult daughters (but not sons) of HSD dams (Tobiansky *et al*. 2021). Interestingly, following mild food restriction, blood corticosterone levels were elevated in both adult daughters and sons of HSD dams (Tobiansky *et al*. 2021). Hence, the HPA axes of daughters and sons are programmed by maternal diet; however, a second “hit” is needed to reveal the change in the HPA axis of sons (Harrell *et al*. 2018).

### Maternal HSD increases aldosterone levels

Aldosterone was significantly increased in specific fetal brain regions (mPFC and NAc) by maternal HSD. This increase of aldosterone, specifically in the mesocorticolimbic system, is another mechanism that could program the brain’s reward circuit. Few studies measure aldosterone when it comes to the effects of stress, even though aldosterone is produced in response to adrenocorticotropic hormone (ACTH). In addition, there are established connections between depression and an increase in the aldosterone/cortisol ratio (hyperaldosteronism) in humans (Murck *et al*. 2023). Aldosterone in the fetus is mainly produced by fetal adrenals, starting at E16 in the rat (Brochu *et al*. 1997; Wotus *et al*. 1998). In humans, circulating aldosterone levels are two to 12-fold higher in the fetus than in the mother (Bayard *et al*. 1970). Bayard et al. also show that maternal and fetal aldosterone can cross the placenta. Pregnancy has no effect on the metabolic clearance rate of aldosterone, but pregnancy increases the rate of aldosterone production (Bayard *et al*. 1970). Here, aldosterone levels are similar in fetal blood and fetal brain, and it is likely that aldosterone in the fetal brain is primarily derived from fetal blood. Nonetheless, E18 rat brain and hippocampal cultures express CYP11B1 and CYP11B2 (MacKenzie *et al*. 2002). Both enzymes are also expressed in adult rat hippocampus and cerebellum (MacKenzie *et al*. 2000). The local upregulation of aldosterone in the mPFC and NAc by maternal HSD could be the result, in part, of local aldosterone synthesis. Aldosterone acts on microglia and reduces their pro-inflammatory activity (Bast *et al*. 2018). As microglia play important roles in brain development, such as guiding neuron migration, synapse formation, or myelination (Gildawie *et al*. 2020; Bordt *et al*. 2024), multiple effects of aldosterone on the mesocorticolimbic system are possible. Future studies should address alterations in microglia morphology or numbers following maternal HSD.

Aldosterone was also increased in other fetal tissues by maternal HSD. For example, sucrose intake significantly increased aldosterone in the labyrinth zone of the placenta, in the blood and in the amniotic fluid of male and female fetuses. Aldosterone also tended to be increased in the maternal serum. Classically, aldosterone is known for regulating electrolyte and fluid balance; however, aldosterone also plays a role in stress physiology (Connell & Davies 2005; Kubzansky & Adler 2010). Moreover, aldosterone has been suggested to act as the main adrenal “stress hormone” in the perinatal period in the rat (Varga *et al*. 2013). During the stress hyporesponsive period (P4-14) in rats (Schmidt 2019), circulating corticosterone is very low at baseline and shows a reduced response to various stressors (Taves *et al*. 2015; Salehzadeh *et al*. 2022). Following a stressor, newborn rat pups show greater changes in aldosterone than corticosterone, in contrast to adult rats (Varga *et al*. 2013). Little is known about the role of aldosterone in fetal development, yet the increases in aldosterone in fetal tissues suggest that maternal sucrose intake is a stressor for the fetus.

### Maternal HSD reduces androgen levels

Androgens were reduced in the placenta by the maternal HSD. In the junctional zone, males and females showed similarly high levels of androstenedione and testosterone. In contrast, in all fetal tissues and the fetus-derived labyrinth zone, males had significantly higher androgen levels than females. During pregnancy, the placenta is an active steroidogenic organ that produces large amounts of progesterone and androgens (Chan & Leathem 1975; Villee 1979; Matt & Macdonald 1984). Interestingly, progesterone, which is required for a healthy pregnancy, tended to be reduced in the junctional zone by maternal HSD. Testosterone levels are particularly high in the junctional zone (Matt & Macdonald 1985). Testosterone is produced by the placenta and serves a critical role in placental development (Parsons & Bouma 2021). The placenta contains androgen receptors, which impact junctional zone formation and function (Gibson *et al*. 2016; Kumar *et al*. 2018; Parsons & Bouma 2021). Testosterone treatment in rats from E14 to E18 increases the size of the junctional zone at E21 (Furukawa *et al*. 2022). Here, maternal HSD reduces placental androgens, which might explain how maternal HSD reduces placenta mass. In one study, maternal aldosterone levels are positively correlated with placental growth, and impaired mineralocorticoid receptor activation is implicated in placenta dysfunction in preeclampsia (Gennari-Moser *et al*. 2011). Here, maternal HSD increases aldosterone levels, and thus changes in aldosterone might not explain the effect of maternal HSD on placenta mass. Glucocorticoids counteract the growth-promoting effect of aldosterone on the placenta (Gennari-Moser *et al*. 2011). Maternal HSD increases corticosterone metabolites and precursors in the dam and fetus (but not placenta). However, the DHC/corticosterone ratio tended to be increased in the junctional zone. In summary, changes in androgens and glucocorticoids might mediate the effect of maternal HSD on placental size, and this is a critical issue for future work.

Maternal HSD significantly decreased androstenedione in the junctional zone and testosterone in the junctional zone and labyrinth zone. These effects are correlated with a reduction in placenta mass, consistent with the idea that local androgen signaling promotes placental growth. Few researchers studying androgens measure androstenedione, and most focus on testosterone. Moreover, microdissection of the placenta is rarely performed but was another useful feature of our experimental design. Follow-up studies should examine androgen receptors and steroidogenic enzymes in the placenta.

Maternal HSD also reduced testosterone in the NAc of the mesocorticolimbic system in the fetal brain. Androgens regulate the expression of tyrosine hydroxylase (TH), a key enzyme in dopamine synthesis, in the mesocorticolimbic system (Adler *et al*. 1999; Kritzer 2003; Wallin-Miller *et al*. 2016; Tomm *et al*. 2022; Seib *et al*. 2023). For example, the androgen synthesis inhibitor abiraterone acetate increases TH+ fibers in the mPFC (Tomm *et al*. 2022), whereas testosterone treatment reduces TH+ fibers in the NAc (Wallin-Miller *et al*. 2016). The reduction in testosterone observed here could alter TH expression in the mesocorticolimbic system during development. Importantly, *Cyp17a1* mRNA levels were significantly reduced in the NAc of adult sons of HSD dams, and CYP17A1 is a critical androgenic enzyme (Tobiansky *et al*. 2021). This result fits very well with the reduced testosterone levels in the NAc seen here. Also, the NAc is particularly sensitive to sucrose and shows large increases in dopamine release following sucrose intake (Hajnal & Norgren 2001; Hajnal *et al*. 2004). Changes in CYP17A1 and androgens in the mesocorticolimbic system could explain the effects of maternal HSD on the behavior of sons, including increased dopamine-dependent motivation for sugar and increased preference for high-sucrose diet and high-fat diet (Tobiansky *et al*. 2021).

Maternal high-sucrose intake leads to broad changes in steroids in the dam, placenta, and fetus. These results raise two important questions: First, how does the sucrose content of our high-sucrose diet compare to common “standard” rat diets? Second, which component of sucrose is responsible for these changes, fructose or glucose? With regard to the first question, one of the standard rodent diets, which has been used for the last 50 years, is Ain76A, formulated by the American Institute of Nutrition. Surprisingly, in Ain76A, over 25% of kcal are from sucrose (Lien *et al*. 2001). According to our data, this amount of sucrose will profoundly alter steroid levels, liver lipids, neurochemistry, and behaviour (present study and Tobiansky et al., 2020, 2021). Another standard rodent diet that has been commonly used since the 1990s is Ain93G. This diet contains 10 % kcal sucrose (Reeves 1997; Lien *et al*. 2001). Thus, many labs around the world have used or are using high- or moderate-sucrose diets without intending to do so. This important point is greatly underappreciated.

With regard to the second question, fructose is generally thought to be responsible for the adverse effects of increased sucrose intake. For example, fructose intake increases the expression of 11β-HSD1 in adipocytes, which increases the regeneration of glucocorticoids (Legeza *et al*. 2017). Similarly, long-term high-fructose intake in male rats increases circulating corticosterone levels (Mosili *et al*. 2020). Adult offspring of rat dams given fructose (but not glucose) during gestation and lactation show an increase in circulating corticosterone levels (Munetsuna *et al*. 2019). Similarly, rat maternal fructose (but not glucose) consumption impairs hippocampal function in the offspring (Yamazaki *et al*. 2018). Placental oxidative stress is linked to high fructose levels in an experimental study in mice and an observational study in humans (Asghar *et al*. 2016). Last, maternal high fructose intake in rats reduces placenta size and alters insulin and leptin signaling in postnatal offspring (Vickers *et al*. 2011). Fructose stimulates AMPK (Kinote *et al*. 2012), unlike glucose (Jiang *et al*. 2021). AMPK alters the HPA axis by modulating glucocorticoid receptor signalling and promoting glucocorticoid resistance (Nader *et al*. 2010; Yuan *et al*. 2016). Taken together, the literature suggests that the adverse effects of a high-sucrose diet are caused by an increase in fructose in the diet.

The effects of maternal HSD show some similarities to the effects of maternal stress. For example, maternal stress in rats reduces 11β-HSD1 expression and activity in the placenta (Mairesse *et al*. 2007). In addition, at E21, fetuses of stressed dams show decreased body, adrenal, pancreas, and testis weights, as well as reduced plasma glucose, growth hormone, and ACTH levels (Mairesse *et al*. 2007). In rats, high-fructose corn syrup (HFCS) intake during gestation and lactation alters activities of 11β-HSD1 and 11β-HSD2 in various tissues (i.e., liver, kidney, adrenal glands, muscle, and white adipose tissue). HFCS intake reduces 11β-HSD2 activity in the dam kidney. Furthermore, epigenetic analysis suggests that miR-27a reduces *Hsd11b2* mRNA expression in the offspring kidney. As a result, maternal HFCS intake increases circulating GC levels in offspring, which may be explained by a decrease in kidney 11β-HSD2 activity (Nouchi *et al*. 2023). In summary, there are commonalities in how maternal stress and high sucrose intake affect steroids in the dam, placenta, and offspring.

In conclusion, a maternal high-sucrose diet has widespread effects on maternal, placental, and fetal steroids, including glucocorticoids, aldosterone, and androgens. The observed changes in the placenta and specific regions of the fetal brain could explain the long-lasting effects of maternal HSD on offspring endocrinology and behavior. Future studies will examine how reduced androgen signaling affects placental growth and function. In addition, we plan to examine microglia and TH in the mesocorticolimbic system.

## ACKNOWLEDGEMENTS

We thank George V. Kachkovski, Griffin Rutledge, Cathy Ma, Asmita Poudel, and Hui W. Chen for help with sample collection, preparation, data acquisition and analysis. We thank Melody Salehzadeh for help with Figure 4, and we thank Marwa Idrissi for help with Figure S1. We thank Dr. Jonathan Spears for help with H&E staining of the placenta.

