## Supplementary file for "Maternal sucrose consumption alters steroid levels in the mother, placenta, and fetus"

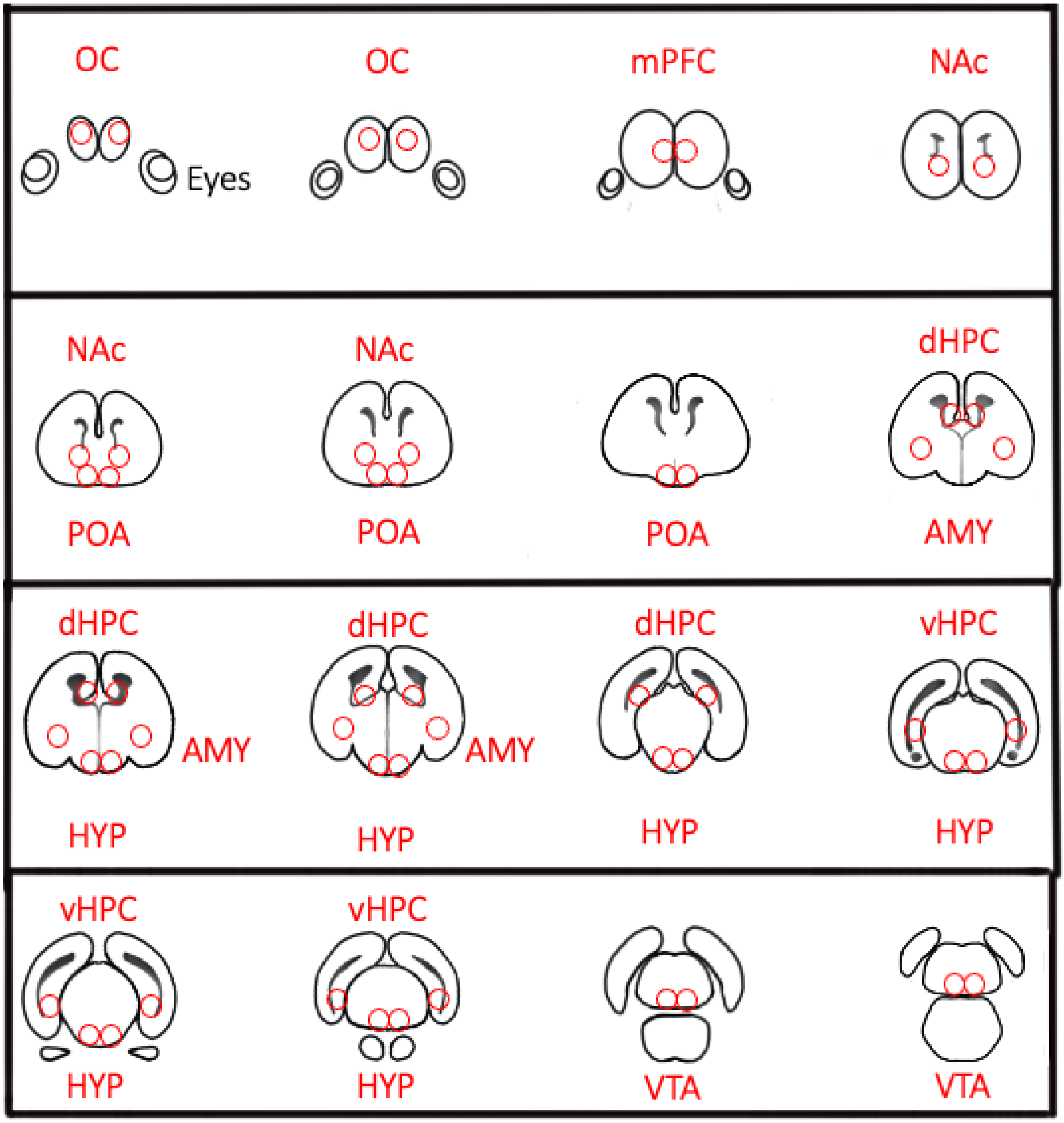


**Figure S1.** Schematic illustrating rostral to caudal locations for microdissection performed using 1 mm diameter Palkovits punch for LC-MS/MS steroid analysis. Section thickness was 300µm. OC, orbital cortex; mPFC, medial prefrontal cortex; NAc, nucleus accumbens; POA, preoptic area; AMY, amygdala; dHPC, dorsal hippocampus; vHPC, ventral hippocampus; HYP, hypothalamus; VTA, ventral tegmental area.

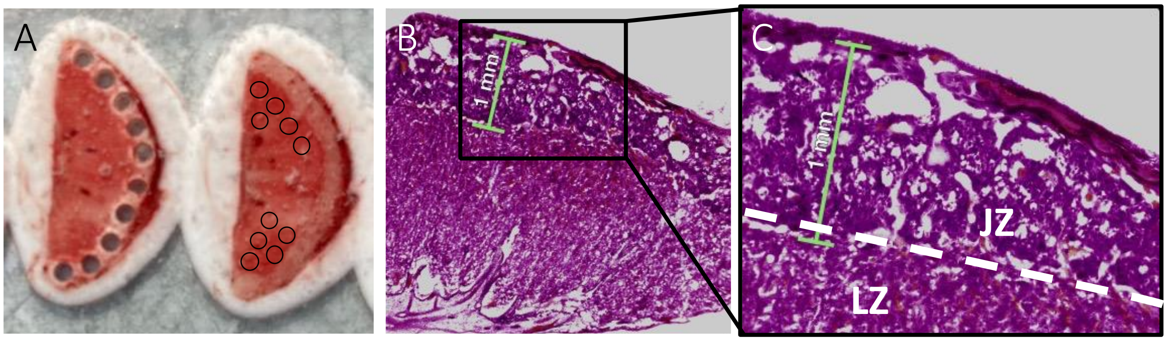


**Figure S2. Punch locations for placenta.** A, Representative 300µm-thick frozen placenta sections. In the section on the left, 1mm-diameter punches of the junctional zone have been collected. In the section on the right, black circles indicate punch locations for the labyrinth zone. B and C, 30µm-thick H&E stained placenta section, highlighting the junctional zone that is 1mm in height. LZ, labyrinth zone; JZ, junctional zone.

Table S1. Steroids (ng/g) in maternal serum that were not significantly affected by HSD

| | CON | HSD |
|---|---|---|
| *Maternal serum* | | |
| Pregnenolone | 4.31 ± 0.62 | 5.89 ± 1.73 |
| Progesterone | 90.97 ± 9.30 | 102.03 ± 3.89 |
| Allopregnanolone | 11.48 ± 1.47 | 13.82 ± 1.06 |
| Androstenedione | 0.76 ± 0.09 | 0.77 ± 0.05 |
| Estrone | 0.02 ± 0.004 | 0.03 ± 0.005 |
| 17β-estradiol | nd | nd |
| Estriol | nd | nd |

Data are shown as mean ± s.e.m. n=10-12. nd, non-detectable. All p-values > 0.05.

Table S2. Steroids (ng/g) in placenta that were not significantly affected by maternal HSD

| | CON Female | | | HSD Female | | | CON Male | | | HSD Male | | |
|---|---|---|---|---|---|---|---|---|---|---|---|---|
| ***Pregnenolone*** | | | | | | | | | | | | |
| Junctional zone | 31.27 | ± | 8.75 | 29.55 | ± | 4.32 | 31.35 | ± | 5.51 | 23.15 | ± | 3.21 |
| Labyrinth zone | 24.22 | ± | 3.36 | 22.00 | ± | 3.27 | 23.11 | ± | 0.92 | 21.88 | ± | 2.19 |
| ***Progesterone*** | | | | | | | | | | | | |
| Junctional zone | **29.43** | **±** | **4.26** | **21.74** | **±** | **3.06** | **25.58** | **±** | **3.37** | **20.71** | **±** | **1.75** |
| Labyrinth zone | 29.48 | ± | 4.24 | 22.63 | ± | 4.61 | 29.73 | ± | 4.40 | 27.81 | ± | 3.50 |
| ***Allopregnanolone*** | | | | | | | | | | | | |
| Junctional zone | 65.61 | ± | 10.47 | 82.23 | ± | 15.43 | 88.67 | ± | 12.79 | 79.39 | ± | 12.16 |
| Labyrinth zone | 76.74 | ± | 15.34 | 89.27 | ± | 16.39 | 118.54 | ± | 19.27 | 74.76 | ± | 16.10 |
| ***Corticosterone*** | | | | | | | | | | | | |
| Junctional zone | 36.02 | ± | 4.83 | 31.51 | ± | 4.68 | 44.34 | ± | 10.28 | 31.60 | ± | 6.79 |
| Labyrinth zone | 54.56 | ± | 6.86 | 53.58 | ± | 9.64 | 80.05 | ± | 18.34 | 52.06 | ± | 7.88 |
| ***11-Dehydrocorticosterone*** | | | | | | | | | | | | |
| Junctional zone | 63.14 | ± | 5.66 | 75.95 | ± | 10.51 | 68.06 | ± | 12.02 | 72.19 | ± | 11.32 |
| Labyrinth zone | 102.91 | ± | 12.40 | 105.61 | ± | 15.19 | 114.18 | ± | 16.73 | 121.00 | ± | 13.47 |
| ***Estrone*** | | | | | | | | | | | | |
| Junctional zone | 0.10 | ± | 0.04 | 0.09 | ± | 0.02 | 0.06 | ± | 0.03 | 0.10 | ± | 0.03 |
| Labyrinth zone | 0.30 | ± | 0.06 | 0.35 | ± | 0.07 | 0.37 | ± | 0.06 | 0.39 | ± | 0.06 |

Data are shown as mean **±** s.e.m. n=6-10. Progesterone showed a trend to be reduced by maternal HSD in the junctional zone (p=0.07), shown in **bold**. All p-values > 0.05. No significant effects of sex. 17β-estradiol and estriol were non-detectable.

Table S3. Steroids (ng/g) in fetal blood and amniotic fluid that were not significantly affected by maternal HSD

| | CON Female | HSD Female | CON Male | HSD Male |
|---|---|---|---|---|
| *Fetal blood* | | | | |
| Pregnenolone | 5.20 ± 1.17 | 4.40 ± 1.25 | 4.28 ± 0.93 | 4.32 ± 0.85 |
| Progesterone | 9.60 ± 0.93 | 8.17 ± 0.74 | 7.92 ± 0.62 | 8.99 ± 0.80 |
| Allopregnanolone | 15.11 ± 3.37 | 18.85 ± 1.90 | 16.82 ± 2.48 | 15.25 ± 1.43 |
| Corticosterone | 192.65 ± 12.89 | 193.72 ± 9.31 | 177.42 ± 15.51 | 171.78 ± 16.00 |
| Estrone | 2.18 ± 0.27 | 2.53 ± 0.15 | 2.21 ± 0.17 | 2.48 ± 0.23 |
| 17β-estradiol | nd | nd | nd | nd |
| Estriol | nd | nd | nd | nd |
| *Amniotic fluid* | | | | |
| Pregnenolone | 0.50 ± 0.12 | 0.67 ± 0.16 | 0.83 ± 0.19 | 0.50 ± 0.07 |
| Progesterone | 0.93 ± 0.08 | 0.92 ± 0.14 | 1.14 ± 0.14 | 1.11 ± 0.18 |
| Allopregnanolone | 2.74 ± 0.22 | 2.94 ± 0.46 | 3.79 ± 0.80 | 3.12 ± 0.28 |
| Corticosterone | 49.86 ± 4.13 | 55.86 ± 7.96 | 60.10 ± 5.46 | 57.58 ± 9.11 |
| Estrone | 0.59 ± 0.05 | 0.68 ± 0.04 | 0.72 ± 0.06 | 0.73 ± 0.05 |
| 17β-estradiol | nd | nd | nd | nd |
| Estriol | nd | nd | nd | nd |

Data are shown as mean ± s.e.m. n=9-11. nd, non-detectable. All p-values > 0.05. No significant effects of sex.

Table S4. Steroids (ng/g) in fetal brain regions

| | CON Female | | | HSD Female | | | CON Male | | | HSD Male | | |
|---|---|---|---|---|---|---|---|---|---|---|---|---|
| *Pregnenolone* | | | | | | | | | | | | |
| OC | 13.60 | ± | 0.74 | 12.49 | ± | 0.94 | 12.24 | ± | 0.71 | 15.04 | ± | 1.56 |
| mPFC | 12.83 | ± | 1.38 | 14.60 | ± | 3.25 | 13.57 | ± | 1.46 | 15.30 | ± | 2.17 |
| NAc | 13.53 | ± | 0.84 | 13.61 | ± | 1.63 | 13.45 | ± | 1.01 | 15.07 | ± | 1.40 |
| AMY | 13.90 | ± | 1.26 | 12.58 | ± | 0.99 | 14.72 | ± | 1.60 | 15.72 | ± | 2.04 |
| POA | 13.96 | ± | 0.98 | 13.21 | ± | 0.76 | 13.10 | ± | 0.94 | 14.24 | ± | 1.37 |
| dHPC | 13.91 | ± | 1.00 | 15.60 | ± | 1.25 | 14.28 | ± | 1.25 | 15.85 | ± | 1.35 |
| vHPC | 16.10 | ± | 1.67 | 16.26 | ± | 1.28 | 14.75 | ± | 0.96 | 16.75 | ± | 1.92 |
| HYP | 18.68 | ± | 1.72 | 16.41 | ± | 1.06 | 16.34 | ± | 1.08 | 20.32 | ± | 1.93 |
| VTA | 21.71 | ± | 1.76 | 20.93 | ± | 1.30 | 20.87 | ± | 1.23 | 23.49 | ± | 2.58 |
| *Progesterone* | | | | | | | | | | | | |
| OC | 27.98 | ± | 2.20 | 23.92 | ± | 4.58 | 24.51 | ± | 3.90 | 25.11 | ± | 2.87 |
| mPFC | 25.91 | ± | 2.68 | 25.93 | ± | 5.35 | 25.85 | ± | 2.59 | 26.10 | ± | 3.62 |
| NAc | 26.85 | ± | 2.42 | 30.29 | ± | 7.58 | 26.76 | ± | 3.88 | 27.47 | ± | 2.71 |
| AMY | 26.12 | ± | 1.69 | 23.72 | ± | 3.86 | 26.44 | ± | 4.60 | 24.67 | ± | 3.00 |
| POA | 26.59 | ± | 2.50 | 25.97 | ± | 2.93 | 26.80 | ± | 3.87 | 25.18 | ± | 2.79 |
| dHPC | 28.06 | ± | 1.98 | 29.22 | ± | 5.88 | 28.15 | ± | 4.71 | 24.57 | ± | 2.31 |
| vHPC | 27.04 | ± | 2.38 | 30.02 | ± | 5.60 | 26.98 | ± | 4.17 | 25.28 | ± | 2.48 |
| HYP | 25.68 | ± | 2.55 | 26.97 | ± | 6.08 | 26.29 | ± | 5.00 | 25.43 | ± | 3.25 |
| VTA | 27.51 | ± | 2.96 | 25.80 | ± | 4.43 | 28.28 | ± | 5.34 | 25.03 | ± | 3.98 |
| *Allopregnanolone* | | | | | | | | | | | | |
| OC | 64.10 | ± | 16.41 | 75.68 | ± | 29.68 | 56.26 | ± | 6.77 | 50.05 | ± | 6.60 |
| mPFC | 67.43 | ± | 13.37 | 81.46 | ± | 20.46 | 81.53 | ± | 13.99 | 54.39 | ± | 10.46 |
| NAc | 51.06 | ± | 11.82 | 64.16 | ± | 16.28 | 56.44 | ± | 6.70 | 45.44 | ± | 5.40 |
| AMY | 45.68 | ± | 9.69 | 59.12 | ± | 18.36 | 52.88 | ± | 8.48 | 42.08 | ± | 5.18 |
| POA | 50.33 | ± | 8.27 | 67.91 | ± | 15.75 | 62.25 | ± | 7.82 | 47.51 | ± | 7.20 |
| dHPC | 40.55 | ± | 5.40 | 50.63 | ± | 10.52 | 48.64 | ± | 6.98 | 36.39 | ± | 4.06 |
| vHPC | 45.73 | ± | 7.50 | 56.88 | ± | 15.12 | 56.54 | ± | 9.71 | 43.88 | ± | 4.49 |
| HYP | 41.38 | ± | 4.05 | 49.53 | ± | 6.23 | 50.82 | ± | 6.84 | 44.37 | ± | 3.51 |
| VTA | 54.78 | ± | 8.54 | 71.65 | ± | 15.60 | 67.85 | ± | 7.39 | 60.15 | ± | 6.05 |

| | CON Female | | | HSD Female | | | CON Male | | | HSD Male | | |
|---|---|---|---|---|---|---|---|---|---|---|---|---|
| *Corticosterone* | | | | | | | | | | | | |
| OC | **30.51** | **±** | **1.71** | **22.98** | **±** | **2.72** | **29.20** | **±** | **5.39** | **24.41** | **±** | **2.05** |
| mPFC | 29.81 | ± | 3.01 | 30.52 | ± | 6.86 | 29.65 | ± | 4.05 | 22.57 | ± | 3.70 |
| NAc | 18.37 | ± | 1.28 | 17.07 | ± | 2.88 | 19.99 | ± | 2.95 | 15.58 | ± | 1.89 |
| AMY | 24.41 | ± | 2.12 | 19.50 | ± | 2.47 | 21.95 | ± | 3.01 | 19.73 | ± | 2.41 |
| POA | 22.35 | ± | 2.19 | 20.64 | ± | 4.30 | 27.44 | ± | 3.92 | 16.51 | ± | 2.63 |
| dHPC | 39.13 | ± | 3.31 | 35.67 | ± | 4.65 | 42.92 | ± | 4.90 | 34.05 | ± | 3.97 |
| vHPC | 27.61 | ± | 2.89 | 29.47 | ± | 4.00 | 30.60 | ± | 4.32 | 27.45 | ± | 2.30 |
| HYP | 32.83 | ± | 2.59 | 28.20 | ± | 3.55 | 30.83 | ± | 5.04 | 24.67 | ± | 3.48 |
| VTA | 27.48 | ± | 2.65 | 28.24 | ± | 4.00 | 37.43 | ± | 6.12 | 26.40 | ± | 2.95 |
| *11-Dehydrocorticosterone* | | | | | | | | | | | | |
| OC | 25.67 | ± | 2.13 | 21.37 | ± | 2.55 | 21.75 | ± | 2.13 | 25.47 | ± | 1.97 |
| mPFC | 28.21 | ± | 2.63 | 30.97 | ± | 5.92 | 28.08 | ± | 2.30 | 27.90 | ± | 2.85 |
| NAc | 32.41 | ± | 3.17 | 30.55 | ± | 2.35 | 27.20 | ± | 2.70 | 32.42 | ± | 3.23 |
| AMY | 30.44 | ± | 2.96 | 29.18 | ± | 3.72 | 24.82 | ± | 1.85 | 29.99 | ± | 2.96 |
| POA | 38.42 | ± | 4.61 | 37.26 | ± | 4.89 | 35.69 | ± | 3.72 | 35.51 | ± | 3.73 |
| dHPC | 31.51 | ± | 3.08 | 33.51 | ± | 3.18 | 32.02 | ± | 3.15 | 31.37 | ± | 2.91 |
| vHPC | 26.82 | ± | 3.00 | 37.84 | ± | 5.11 | 28.62 | ± | 1.59 | 31.11 | ± | 2.15 |
| HYP | 31.42 | ± | 3.68 | 33.06 | ± | 4.17 | 28.02 | ± | 2.70 | 31.32 | ± | 3.14 |
| VTA | 29.23 | ± | 2.55 | 33.61 | ± | 4.24 | 29.06 | ± | 2.82 | 32.07 | ± | 3.75 |
| *Aldosterone* | | | | | | | | | | | | |
| OC | 0.07 | ± | 0.01 | 0.12 | ± | 0.05 | 0.06 | ± | 0.01 | 0.21 | ± | 0.08 |
| mPFC | **0.06** | **±** | **0.003** | **0.2** | **±** | **0.04** | **0.06** | **±** | **0.01** | **0.12** | **±** | **0.03** |
| NAc | **0.08** | **±** | **0.01** | **0.21** | **±** | **0.08** | **0.09** | **±** | **0.02** | **0.21** | **±** | **0.08** |
| AMY | 0.08 | ± | 0.018 | 0.19 | ± | 0.054 | 0.12 | ± | 0.01 | 0.21 | ± | 0.086 |
| POA | 0.08 | ± | 0.01 | 0.17 | ± | 0.06 | 0.08 | ± | 0.01 | 0.23 | ± | 0.11 |
| dHPC | 0.11 | ± | 0.03 | 0.24 | ± | 0.09 | 0.14 | ± | 0.027 | 0.28 | ± | 0.11 |
| vHPC | 0.09 | ± | 0.02 | 0.25 | ± | 0.10 | 0.12 | ± | 0.03 | 0.26 | ± | 0.10 |
| HYP | 0.12 | ± | 0.02 | 0.22 | ± | 0.08 | 0.13 | ± | 0.02 | 0.24 | ± | 0.09 |
| VTA | 0.09 | ± | 0.02 | 0.24 | ± | 0.10 | 0.12 | ± | 0.028 | 0.29 | ± | 0.108 |

| | CON Female | | | HSD Female | | | CON Male | | | HSD Male | | |
|---|---|---|---|---|---|---|---|---|---|---|---|---|
| *Androstenedione* | | | | | | | | | | | | |
| OC | 1.29 | ± | 0.21 | 0.96 | ± | 0.17 | 1.83 | ± | 0.29 | 1.96 | ± | 0.16 |
| mPFC | 1.40 | ± | 0.32 | 0.97 | ± | 0.12 | 2.22 | ± | 0.37 | 2.20 | ± | 0.19 |
| NAc | 1.21 | ± | 0.19 | 0.92 | ± | 0.16 | 1.88 | ± | 0.34 | 1.95 | ± | 0.16 |
| AMY | 0.89 | ± | 0.14 | 0.72 | ± | 0.13 | 1.38 | ± | 0.32 | 1.60 | ± | 0.19 |
| POA | 1.01 | ± | 0.21 | 0.72 | ± | 0.15 | 1.50 | ± | 0.22 | 1.35 | ± | 0.19 |
| dHPC | 1.20 | ± | 0.14 | 0.88 | ± | 0.13 | 1.85 | ± | 0.28 | 1.80 | ± | 0.18 |
| vHPC | 1.27 | ± | 0.20 | 1.02 | ± | 0.13 | 2.04 | ± | 0.31 | 1.99 | ± | 0.18 |
| HYP | 1.21 | ± | 0.23 | 0.80 | ± | 0.12 | 1.64 | ± | 0.21 | 1.89 | ± | 0.20 |
| VTA | 1.41 | ± | 0.25 | 1.04 | ± | 0.11 | 2.05 | ± | 0.32 | 2.27 | ± | 0.24 |
| *Testosterone* | | | | | | | | | | | | |
| OC | 0.06 | ± | 0.011 | 0.04 | ± | 0.01 | 1.2 | ± | 0.32 | 0.97 | ± | 0.11 |
| mPFC | 0.05 | ± | 0.003 | 0.05 | ± | 0.002 | 1.2 | ± | 0.23 | 1.0 | ± | 0.15 |
| NAc | **0.08** | **±** | **0.02** | **0.03** | **±** | **0.007** | **1.2** | **±** | **0.28** | **1.0** | **±** | **0.1** |
| AMY | 0.05 | ± | 0.008 | 0.03 | ± | 0.01 | 1.1 | ± | 0.35 | 0.92 | ± | 0.11 |
| POA | 0.04 | ± | 0.01 | 0.03 | ± | 0.01 | 1.3 | ± | 0.28 | 1.0 | ± | 0.13 |
| dHPC | 0.06 | ± | 0.007 | 0.04 | ± | 0.006 | 1.3 | ± | 0.37 | 1.0 | ± | 0.1 |
| vHPC | 0.05 | ± | 0.006 | 0.05 | ± | 0.008 | 1.3 | ± | 0.36 | 1.1 | ± | 1.1 |
| HYP | 0.06 | ± | 0.005 | 0.04 | ± | 0.007 | 1.3 | ± | 0.3 | 1.1 | ± | 0.13 |
| VTA | 0.05 | ± | 0.006 | 0.03 | ± | 0.005 | 1.4 | ± | 0.39 | 1.3 | ± | 0.18 |
| *Estrone* | | | | | | | | | | | | |
| OC | 0.27 | ± | 0.03 | 0.27 | ± | 0.03 | 0.27 | ± | 0.06 | 0.30 | ± | 0.03 |
| mPFC | 0.27 | ± | 0.04 | 0.27 | ± | 0.06 | 0.29 | ± | 0.09 | 0.29 | ± | 0.03 |
| NAc | 0.35 | ± | 0.04 | 0.31 | ± | 0.03 | 0.33 | ± | 0.08 | 0.37 | ± | 0.04 |
| AMY | 0.60 | ± | 0.06 | 0.55 | ± | 0.09 | 0.60 | ± | 0.08 | 0.72 | ± | 0.10 |
| POA | 0.89 | ± | 0.11 | 0.84 | ± | 0.09 | 1.12 | ± | 0.08 | 1.27 | ± | 0.10 |
| dHPC | 0.43 | ± | 0.04 | 0.42 | ± | 0.06 | 0.47 | ± | 0.05 | 0.47 | ± | 0.04 |
| vHPC | 0.28 | ± | 0.03 | 0.36 | ± | 0.04 | 0.34 | ± | 0.04 | 0.37 | ± | 0.04 |
| HYP | 0.55 | ± | 0.07 | 0.52 | ± | 0.07 | 0.52 | ± | 0.03 | 0.64 | ± | 0.04 |
| VTA | 0.25 | ± | 0.03 | 0.29 | ± | 0.04 | 0.37 | ± | 0.03 | 0.36 | ± | 0.04 |

Data are shown as mean **±** s.e.m. n=6-10. For 17β-estradiol and estriol, less than 20% of samples had detectable levels. Two-way ANOVA was used for statistical analysis to examine the effects of diet and sex on steroids in different brain regions. No diet x sex interactions were observed. Androstenedione and testosterone were significantly higher in male fetuses compared to female fetuses across all brain regions. Main effect of diet: data shown in **bold**, $p \leq 0.05$. OC, orbital cortex; mPFC, medial prefrontal cortex; NAc, nucleus accumbens; AMY, amygdala; POA, preoptic area; dHPC, dorsal hippocampus; vHPC, ventral hippocampus; HYP, hypothalamus; VTA, ventral tegmental area; CON, control; HSD, high-sucrose diet.
